# Non-additive dosage-dependent effects of *TaGS3* gene editing on grain size and weight in wheat

**DOI:** 10.1101/2024.04.28.591550

**Authors:** Wei Wang, Qianli Pan, Bin Tian, Dwight Davidson, Guihua Bai, Alina Akhunova, Harold Trick, Eduard Akhunov

## Abstract

The grain size in cereals is one of the main component traits contributing to yield. Previous studies showed that loss-of-function (LOF) mutations in *GS3*, encoding Gγ subunit of the multimeric G protein complex, increase grain size and weight in rice. While association between allelic variation in *GS3* homologs of wheat and grain weight/size was detected previusly, the effects of LOF alleles on these traits remain unknown. We used genome editing to create the *TaGS3* mutant lines with the LOF homeo-allele dosage variation. Contrary to results obtained for rice, editing of all three *TaGS3* copies result in significant decrease in grain length, width, grain area and weight, without affecting number of grains per spike. Compared to wild type, the highest increase in grain weight and area was observed in mutants with the intermediate dosage of the LOF alleles, indicating that suppressive effects of *TaGS3* on grain size and weight in wheat are dosage-dependent and non-additive. Our results suggest that *TaGS3* likely represents a functionally diverged homolog of *GS3* evolved in the wheat lineage. The newly developed LOF alleles of *TaGS3* expand the set of CRISPR-Cas9-induced variants of yield component genes that could be used for increasing grain weight in wheat.

## Introduction

Genetic analyses of yield component traits, mostly performed in rice, identified a number of genes involved in pathways controlling grain size and weight in cereal crops (Li and Li 2016; Li and Yang 2017). Among many of the characterized grain size/weight QTL loci identified in rice are *GW2* (Song *et al*. 2007), *GS3* (Fan *et al*. 2006), *GW8* (Wang *et al*. 2012), *GW7* (Wang *et al*. 2015), *GS5* (Li *et al*. 2011), *GS2* (Hu *et al*. 2015), *GSE5* (Geng *et al*. 2017), *OsCKX2* (Ashikari *et al*. 2005), *DEP1* (Huang *et al*. 2009) and *GS3* (Fan *et al*. 2006). identification of these genes along with the development of comparative genomic resources accelerated genetic dissection of yield component traits in other crops, including wheat. The analysis of natural allelic diversity and/or mutagenesis performed in wheat showed that many rice homologs of grain size and weight QTL are linked with variation in the same traits. For example, both gene editing and mutagenesis showed that the loss-of-function mutations in *TaGW2* gene encoding RING-type protein with E3 ubiquitin ligase result in increased grain size and weight (Simmonds *et al*. 2016; Wang *et al*. 2018b). The editing of *TaGW7* gene encoding TONNEAU1-recruiting motif protein demonstrated its involvement in regulation of grain shape and weight in wheat (Wang *et al*. 2019). Likewise, natural variation in the *TaCKX2* (Zhang *et al*. 2012) and *TaGS3* (Yang *et al*. 2019) genes was associated with variation in grain weight traits in wheat.

One of the well investigated pathways contributing to grain size variation is mediated by G protein complex composed of Gα, Gβ and Gγ subunits (Thung *et al*. 2012; Urano and Jones 2014). In rice, *GS3* gene encoding the atypical Gγ subunit was shown to act as negative regulator of grain size and weight (Fan *et al*. 2006). *GS3* competes for interaction with the Gβ subunit with two other atypical Gγ subunits, DEP1 and GGC2, that act as positive regulators of grain size and weight (Sun *et al*. 2018). The N-terminal organ size regulation (OSR) domain of *GS3* was shown to be critical for negative regulatory effects (Mao *et al*. 2010). Loss-of-function mutations in *GS3* resulting in longer grain and increase in grain weight were suggested to remove the repressive effect of *GS3* (Sun *et al*. 2018). The strongest positive impact of grain length in rice was obtained in lines with the knock-out allele of *GS3* and the overexpressed *DEP1* and *GGC2* genes (Sun *et al*. 2018). Allelic variation in *GS3* was shown to substantially contribute to variation in grain size and weight in natural populations and played important role in increasing rice productivity (Mao *et al*. 2010).

In wheat, significant association was found between natural variation in the *TaGS3-4A* and *TaGS3-7A* homoeologs and grain weight and length (Yang *et al*. 2019; Zhang *et al*. 2020), with different splicing variants of the gene having distinct effects on these traits (Ren *et al*. 2021). Interestingly, overexpression of five splicing variants of *TaGS3* in wheat showed that one of the gene isoforms have positive effect on grain size and weight in wheat (Ren *et al*. 2021). It was suggested that alternative splicing resulting in truncation of the OSR domain in this *TaGS3* isoform reduces its affinity to Gβ domain, reducing its negative impact on grain size and weight (Ren *et al*. 2021). These studies suggest that the *TaGS3* gene, like its homolog in rice, should also act as a negative regulator of pathways controlling grain size and weight. Therefore, one might expect that the complete knock-out of *TaGS3* in wheat should have strong positive effect on grain size and weight.

Here, we used CRISPR-Cas9-based gene editing system for creating the LOF mutants of the three homoeologous copies of the *TaGS3* gene in the wheat genome. The effects of gene editing on grain morphometric traits, grain weight and grain number per head was investigated in populations derived from independent transformation events. Contrary to expectations based on prior functional studies of *GS3* in rice and *TaGS3* in wheat, the triple-knockout mutants of *TaGS3* with non-functional alleles in all three wheat genomes showed significant decrease in grain length, width, grain area and weight, without discernable changes in the number of grains per spike. The highest increase in grain weight and area was obtained in lines carrying the intermediate number the LOF alleles, suggesting that suppressive effects of *TaGS3* on grain traits in wheat are dosage-dependent rather than additive. Our results also suggest that *TaGS3* in wheat and its rice homolog *GS3* are not functionally equivalent and that intermediate levels of *TaGS3* expression are likely necessary for optimal grain size and weight trait expression in wheat.

## Results

### CRISPR-Cas9 editing of the *GS3* homologs in wheat

Bread wheat has three copies of a gene showing highest similarity to *GS3* from rice are located on chromosomes 7A (TraesCS7A02G017700), 4A (TraesCS4A02G474000 TraesCS4A02G474000) and 7D (TraesCS7D02G015000). The length of *TaGS3* homoeologs (170, 169, 121, and 169 amino acids) in wheat is shorter than that of rice *GS3* (232 amino acids) due to differences in the length of the cysteine rich tails (Fig. 1a). The cysteine-rich domain was previously shown to play a role in the rate regulation of protein degradation, with longer cysteine-rich domain resulting in faster degradation of *GS3* in rice and reduced repressive effect on grain size (Sun *et al*. 2018). However, the effect of tail length variation on grain size is substantially smaller than the effect of the OSR domain, which plays major role in interaction with the Gβ (Mao *et al*. 2010).

**Fig. 1.**
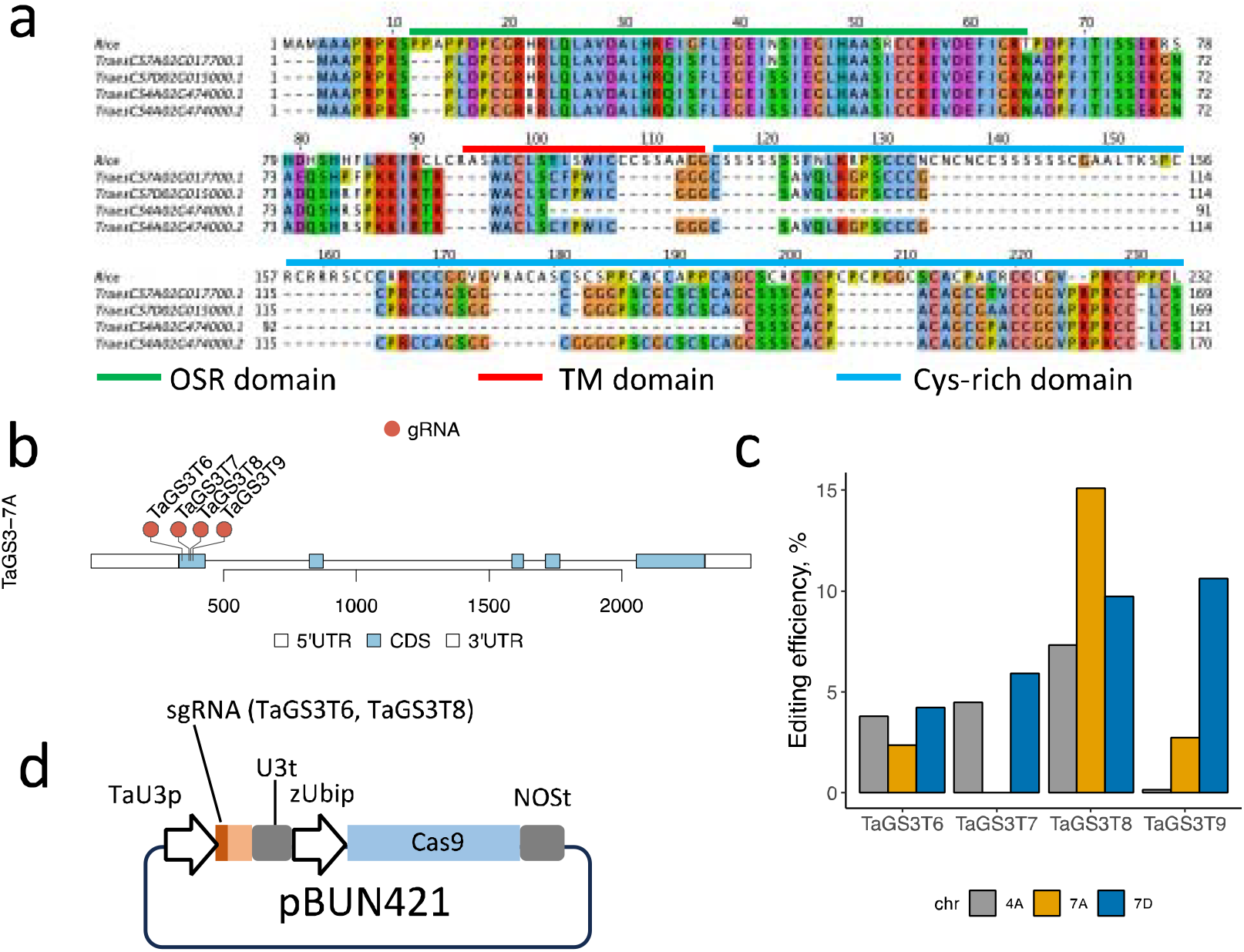
CRISPR-Cas9 editing of the *GS3* homologs in wheat. **A**. Protein sequence alignment of rice and wheat GS3 homologs. The boundaries of functional domains (OSR, trans-membrane (TM) and cysteine-rich) are shown relative to rice GS3 protein sequence (Fan *et al*. 2006). **B**. Distribution of four gRNA target sites in the *TaGS3* gene (*TaGS3-7A* homoeolog is used as example). **C**. Gene editing efficiency assessed in the wheat protoplasts. The proportion of Illumina reads with editing events at the target sites was calculated for each TaGS3 homoeolog by sequencing target sites using DNA extracted from the wheat protoplasts. The values were normalized by the protoplast transformation efficiency assessed using a plasmid expressing the YFP fluorescent protein. **D**. Two pooled gene editing constructs targeting TaGS3T6 and TaGS3T8 sites were used for biolistic transformation of wheat cultivar Bobwhite.

To maximize the functional effect of mutations, we designed four gRNAs targeting the conserved OSR domain coding regions in the first exon of the *TaGS3* homoeologs (Fig. 1b, Supplementary Table 1). The gRNAs have been subcloned into plasmid pBUN421 and their editing efficiency was evaluated by transiently expressing constructs in the wheat protoplasts. The editing efficiency assessed by next-generation sequencing of PCR products including the gRNA target sites (Wang *et al*. 2021) ranged from 0 to 15.1% (Fig. 1c). Based on the efficiency of editing, we selected plasmids with sgRNAs targeting the GS3T6 and GS3T8 sites for biolistic transformation (Fig. 1d). A total 67 independent transgenic plants in cultivar Bobwhite carrying the Cas9-gRNA constructs have been regenerated, and two transgenic lines, 4906-1 and C538-1, have been used for developing populations to evaluate the effects of gene editing in *TaGS3* on yield component traits. Both lines carried mutations resulting in premature termination codons in all three homoeologous copies of *TaGS3*.

### The effect of *TaGS3* gene editing on yield component traits

To reduce the possible effects of epigenetic changes associated with the regeneration of transgenic lines, the transgenic line 4906-1 that carried mutations in all three copies of the *TaGS3* gene was crossed with wild-type cultivar Bobwhite. The BC_1_F_2_ population derived from this cross segregated for the number of functional *TaGS3* gene copies, which allowed us to investigate the effect of allele dosage on yield component traits. The BC_1_F_2_ population was phenotyped for thousand grain weight (TGW), grain length (GL), grain width and grain area (GA) traits (Fig. 2). The lines were further grouped based on the number of non-edited wild-type alleles and the mean phenotypic values were compared between the groups. The main effect of reduction in the number of functional *TaGS3* alleles in this population was the reduction in grain length (Fig. 2a), which is opposite to the effects expected from *TaGS3* editing if this gene is a negative regulator of grain length. The lines that carried five non-functional copies of *TaGS3* showed 2.5% reduction (t-test p-value = 0.02) in GL compared to lines with only one functional copy of *TaGS3*. No significant effects of *TaGS3* editing were detected for other grain morphometric traits (Fig. 2a). In lines that carried intermediate number of wild type alleles, we have also observed some increase in the TGW compared to lines that carried 1 or 5 wild type alleles (Fig. 2a).

**Fig. 2.**
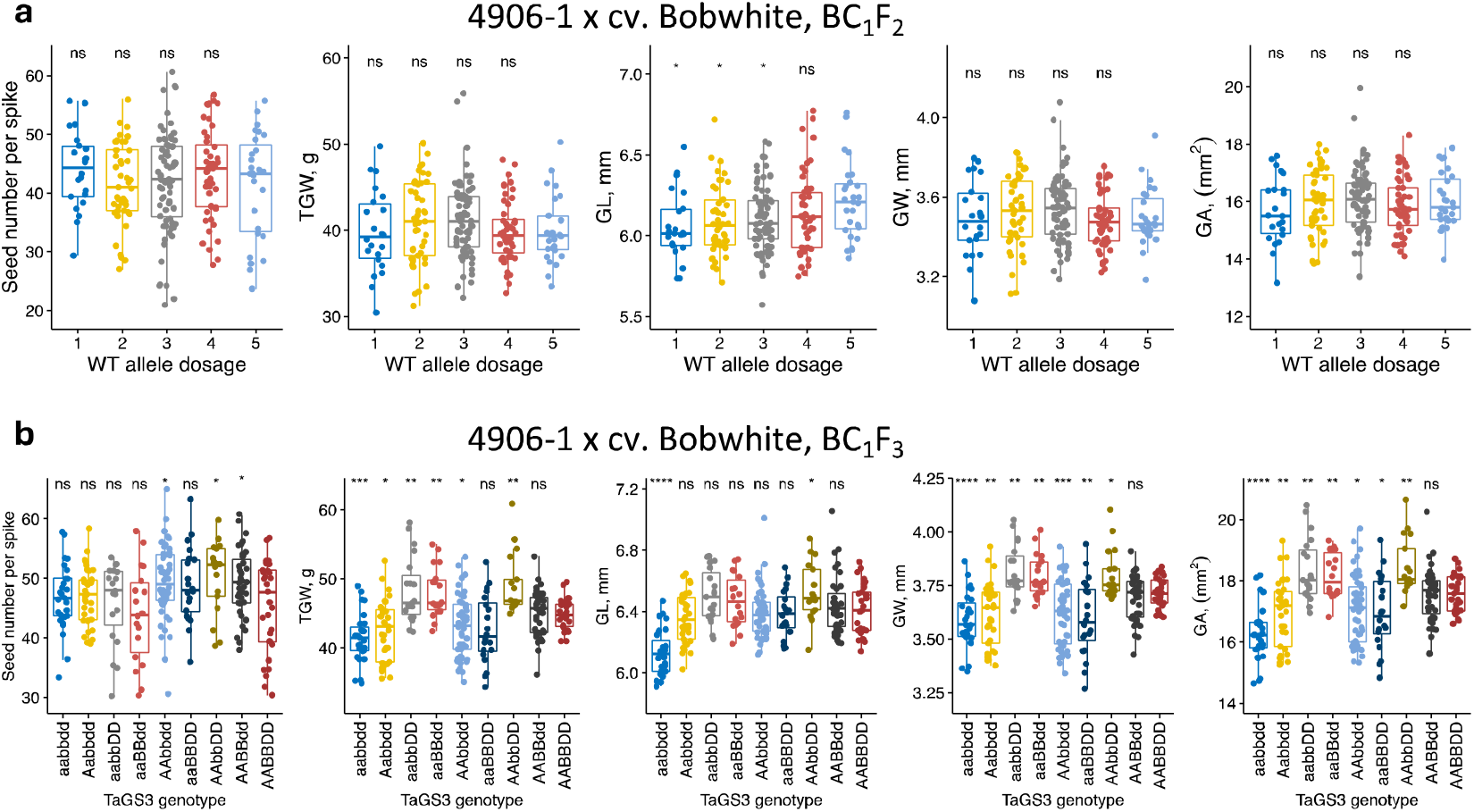
The effects of *TaGS3* gene editing on yield component traits assessed in the BC_1_F_2_ and BC_1_F_3_ populations. **a**. The BC_1_F_2_ population was created by crossing 4906-1 line with wild-type cultivar Bobwhite. The phenotypic measurements were conducted using lines grouped based on the total number of functional wild-type (WT) *TaGS3* alleles at three loci located on chromosomes 7A, 4A and 7D. **b**. The phenotypic measurements were conducted using lines grouped based on their genotypes at three *TaGS3* homoeologous loci. The upper- and lowercase letters correspond to genotypes of the functional (A, B or D) and edited (a, b or d) *TaGS3* alleles, respectively. The graphs show relationships between grain number, morphometric traits and weight and the number of functional wild-type *TaGS3* alleles. The significance testing was performed by applying *t*-test to compare each genotype group with a reference group that includes lines with the highest number of functional wild-type alleles in each population. Significance levels: **** - ≤ 0.0001, *** - ≤ 0.001, ** - ≤ 0.01, * - ≤ 0.05, ns - > 0.05.

To confirm the observed effects, we developed BC_1_F_3_ population using lines from the BC_1_F_2_ population that carried knock-out mutations at one (genotypes aaBBDD, AAbbDD, or AABBdd), two (genotypes aabbDD, AAbbdd, or aaBBdd) or three (aabbdd) *TaGS3* loci (Fig. 2b). In this population, we have also observed reduction in GL with increase in the number of edited gene copies. While single- or double-gene mutants did not show significant differences in GL compared to wild-type genotype, the triple mutants showed 4.4% reduction in GL (t-test p-value = 1.7 x 10^-8^), confirming observations made using the BC_1_F_2_ population.

The trend observed for TGW in the BC_1_F_2_ population, where higher TGW was observed in lines carrying intermediate number of knockout mutations than in lines with all alleles being mutated or wild type, became more pronounced in the BC_1_F_3_ population. The knockouts in all *TaGS3* copies led to 7.5% reduction in TGW relative to wild-type (p-value = 1.5 x 10^-4^). On contrary, in two double-mutants with genotypes aabbDD and aaBBdd, we observed 7.9% (p-value = 6.4 x 10^-3^) and 7.2% (p-value = 2.4 x 10^-3^) increase in TWG, respectively, compared to wild type line. The third double mutant with genotype AAbbdd, however, showed 3.8% reduction in TGW (p-value = 2.5 x 10^-2^). The TGW of two single mutant lines (AABBdd and aaBBDD) showed no significant difference from that of the wild-type line. The only single-locus mutant AAbbDD that showed 9.5% increase in TGW (p-value = 1.1 x 10^-3^) was located in *TaGS3-4A*, which is also the homoeolog that is expressed at the highest level in developing grain (Zhang *et al*. 2020). The relationships observed between the genotypes of mutated *TaGS3* and TGW were also similar to those observed for GW and GA traits (Fig. 2b). Similar to results obtained in the BC_1_F_2_ population, the *TaGS3* gene editing had no detectable effects on grain number per spike.

In addition, the phenotypic effects of *TaGS3* editing were validated in the T_5_ generation population derived from T_0_ transgenic plant C538-1. In T_0_ generation this plant showed low editing frequency and had only two *TaGS3* copies mutated. In T_3_ generation of this line, we have recovered line C538-1-78-2-22, which was heterozygous at all three homoeologous *TaGS3* loci. The gene expression analysis showed that this line also lacks Cas9 expression, likely due to silencing of Cas9 in the previous generations. This gave us an opportunity to develop a population of 400 T_5_ generation lines that segregated at all three *TaGS3* loci (Fig. 3).

**Fig. 3.**
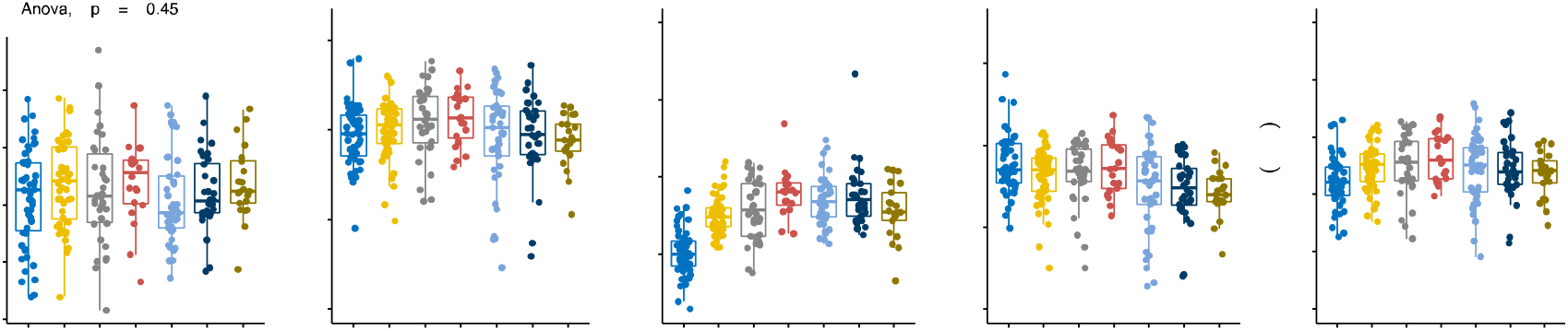
Phenotypic evaluation of *TaGS3* gene editing in the T_5_ generation population. The population was derived from C538-1 transgenic plant. The phenotypic measurements were conducted using lines grouped based on the total number of wild-type (WT) *TaGS3* alleles at three loci located on chromosomes 7A, 4A and 7D. The graphs show relationships between grain number, morphometric traits and weight and the number of functional *TaGS3* alleles. The significance testing was performed by applying post-hoc *t*-test to compare each genotype group with a reference group that includes lines with the highest number of functional wild-type alleles in each population. Significance levels: **** - ≤ 0.0001, *** - ≤ 0.001, ** - ≤ 0.01, * - ≤ 0.05, ns - > 0.05.

Consistent with the results obtained in the populations derived from transgenic line 4906-1 (Fig. 2), the group of lines in T_5_ population with all copies of *TaGS3* edited showed 2.8% reduction in GL compared to wild-type line (p-value = 1.2 x 10^-6^). Similar to results from 4906-1-derived populations, we have also observed non-additive dosage effects for TGW and GA with highest increase observed in lines carrying intermediate number of functional wild-type *TaGS3* alleles (Fig. 4). While no significant difference in TGW and GA was observed between the wheat lines from groups with all copies of *TaGS3* either edited or non-edited, group of lines with three copies of the functional *TaGS3* allele showed 3.4% increase in TGW (p-value = 9.1 x 10^-3^) and 2.1% increase in GA (p-value = 1.9 x 10^-2^) compared to wild type line. Thus, decrease or increase of the number of functional *TaGS3* alleles below or above three, respectively, was associated with decrease in TGW and GA. In T_5_ population, the GW trait showed a trend towards increase with decrease in the number of the functional *TaGS3* alleles. However, this trend was not supported by observations made in the BC_1_F_2_ and BC_1_F_3_ populations (Fig. 2), and likely could be attributed to genotype-specific epigenetic modifications associated with regeneration of C538-1 transgenic plant. In our previous studies, we found that it is important to take this factor into account and perform phenotypic evaluation of transgenic lines in advanced generations after crossing with wild type line (Wang *et al*. 2018b).

Overall, the results of the analyses performed in two advanced generation populations derived from independently developed transgenic plants 4906-1 and C538-1 were consistent for the GL, TGW and GA traits. Our results indicate that 1) fixation of all homoeologous copies of *TaGS3* for LOF alleles results in GL, TGW and GA reduction, 2) optimal expression of GL, TGW and GA traits in edited lines is observed in lines with the intermediate dosage of the LOF *TaGS3* alleles, and 3) certain combination of the LOF homeo-alleles have significant positive effect on GL, TGW and GA traits suggestive of epistatic interaction between genomes.

## Discussion

The analyses of natural variation in the *TaGS3* gene in wheat (Yang *et al*. 2019; Zhang *et al*. 2020; Ren *et al*. 2021) and functional and diversity analyses of *GS3* in rice (Fan *et al*. 2006; Takano-Kai *et al*. 2009; Sun *et al*. 2018) indicate that *GS3* homologs are negative regulators of grain size and weight in both crops. However, effects of LOF in the wheat *TaGS3* homeologs on grain size and weight traits were not investigated. In our study, we show that contrary to results obtained in rice (Sun *et al*. 2018), where the LOF alleles of GS3 are associated with highest increase in grain length and weight, the fixation of *TaGS3* loci from three wheat genomes for LOF allele results in significant decrease in grain length, grain size and weight. These results suggest that *TaGS3* is functionally diverged from its *GS3* rice homolog. The functionality of the *TaGS3* gene homoeologs, at least some of them, appear to be important for optimal expression of grain dimension and weight traits in wheat.

The regulatory effects of *TaGS3* on grain-related traits are more complex in wheat compared to those observed in rice, likely due to the presence of three homoeologous copies. First, we found that the most substantial positive effects on grain size and weight traits were observed in edited lines that carry the intermediate number of LOF *TaGS3* alleles. The effects of the *TaGS3* gene editing were dosage-dependent but non-additive, suggesting that the accumulation of the LOF homeo-alleles have positive effect only to the certain levels and that *TaGS3* expression is still required for optimal expression of grain-related traits in wheat.

Second, the positive effects of the *TaGS3* LOF allele accumulation could be associated with only some of the wheat genomes. In our study, we found that the 7A-4A and 7A-7D combinations of the LOF homeo-alleles have significant positive effect on grain weight and dimensions, whereas 4A-7D LOF allele combination had negative effect. The analyses of natural genetic diversity in wheat associated significant increases in TGW and GL with two *TaGS3* haplotypes on chromosomes 4A and 7D, suggestive of epistatic interaction between these homoeologous copies of genes in wheat (Zhang *et al*. 2020). It is possible that in our populations, 4A-7D LOF combination reduces the positive effect of epistasis between these homeo-loci on grain size and weight.

Third, the results of gene editing indicate that each of the three homoeologous copies of *TaGS3* are functional and have potential to influence grain size and weight traits in wheat. This observation is consistent with the analyses of natural genetic diversity that linked variation in all three *TaGS3* gene copies from chromosomes 7A, 4A and 7D with variation in grain size and weight in wheat (Yang *et al*. 2019; Zhang *et al*. 2020). The differences in the phenotypic impact of editing distinct homoeologous copies of *TaGS3* could potentially be associated with cultivar-and genome-specific differences in expression and alternative splicing. Previously, we have demonstrated that the cultivar-specific phenotypic effects of editing distinct homoeologous copies of *TaGW2* on grain size and weight are associated with the expression levels in different cultivars (Wang *et al*. 2018b). Earlier studies in wheat (Ren *et al*. 2021) and rice (Liu *et al*. 2022) showed that AS plays a role in controlling the proportion of *GS3* isoforms that have different impact on grain size and weight traits. While in rice only two isoforms, both having suppressive OSR domain, were identified (Liu *et al*. 2022), in wheat five AS isoforms were detected for each of the three *TaGS3* homoeologs (Ren *et al*. 2021), with one of the isoforms with truncated OSR domain having strong positive impact on grain size and weight. This observation might potentially explain the negative impact of triple mutations in *TaGS3* on grain size, length and weight and positive impact of 7A-4A and 7A-7D homoeolog editing on these traits found in our study.

Overall, our work provides further insights into the complex genetic control of grain dimension and weight traits by the *TaGS3* homoeologous loci in wheat. Our results indicate that *TaGS3* acts not only as a negative regulator of grain length and weight in wheat. The presence of certain number of the functional *TaGS3* gene copies appears to be critical for optimal trait expression. Using CRISPR-Cas9 editing, we have developed the LOF homeo-alleles for *TaGS3* and identified the single- and two-locus combinations of these alleles that have potential to increase grain length and weight. Combined with alleles identified in natural populations these LOF *TaGS3* alleles could be used for improving yield potential in wheat breeding programs.

## Materials and Methods

### Plant growth conditions

The edited plants were grown in greenhouse under 16-hour light/8-hour dark and the temperature set at 24 °C in the day and 21 °C in the night. The T_1_ generation plants were grown in the 0.2 L square pots filled with SunGro soil (Sun Gro Horticulture, Agawam, MA, USA). All other plants were grown in 1 liter square pots filled on 3/4 with soil mix (volume ratio soil:peatmoss:perlites:CaSO_4_ is 20:20:10:1) and the top ¼ with SunGro soil mix (Sun Gro Horticulture, Agawam, MA, USA). Plants in greenhouse were arranged according to the complete randomized design.

### Analysis of the *TaGS3* sequences

The rice *GS3* gene CDS (Fan *et al*. 2006) was used to perform BLASTN search against the wheat reference genome RefSeq v2.0 (The International Wheat Genome Sequencing Consortium (IWGSC) 2018) in the Ensembl Plants website (plants.ensembl.org). The orthologous genes were identified using the sequence similarity threshold above 70% and located on chromosomes 7A, 4A and 7D syntenic to rice chromosome 3. The annotated sequences of the wheat orthologs, 7A (TraesCS7A02G017700), 4A (TraesCS4A02G474000 TraesCS4A02G474000) and 7D (TraesCS7D02G015000), were downloaded from Ensembl Plants and henceforth, will be referred to as *TaGS3-7A, TaGS3-4A* and *TaGS3-7D*, respectively. The protein sequences of genes including both introns and exons were used to build phylogenetic tree. Alignment was performed using MUSCLE (Edgar 2004).

### Design and validation of CRISPR-Cas9 targets on *TaGS3* gene

The CIRSPR-Cas9 targets on *TaGW3* gene were designed as described previously (Wang *et al*. 2018b, 2019). The coding regions of *TaGS3* homoeologs were aligned to identified conserved sequences, which were analyzed using sgRNAscorer 1.0 (http://crispr.med.harvard.edu/sgRNAScorer). The top ranked targets were compared using BLASTN against the wheat genome RefSeq v2.0 (The International Wheat Genome Sequencing Consortium (IWGSC) 2018) to select targets with low potential for off-target editing. A total four gRNAs were designed to target the region near the CDS start in the first exon. The sequence of each gRNA was synthesized as two complementary oligos with 4 nucleotides as overhangs at both termini (Supplementary Table 1). The oligos were annealed and sub-cloned into CRISPR-Cas9 plasmid pBUN421 as described (Wang *et al*. 2018a). The genome editing efficiencies of the designed gRNAs were estimated by transiently expressing the constructs in the wheat protoplasts followed by the amplification and next generation sequencing (NGS) of target flanking sequences (Wang *et al*. 2019).

### Regeneration of transgenic plants and genotyping of mutants

The constructs targeting two target sites TaGW3T6 and TaGW3T8 were mixed in equimolar amounts with the *bar*-gene carrying pAHC20 construct and ballistically transformed into wheat embryos. The T0 transgenic plants were regenerated as in our previous study (Wang *et al*. 2019). Three primer pairs spanning the CRISPR-Cas9 constructs were used to screen for Cas9 positive plants by PCR (Table S1). The target sites of the CRISPR-Cas9 positive plants and their T1, T2 and T3 progenies were genotyped by the NGS of pooled barcoded PCR amplicons, as described previously (Wang *et al*. 2018a). The primers used for genotyping are included in the Supplementary Table 1.

### Collection of grain dimension, grain number per spike and TGW (thousand grain weight) data

The grain size (grain width, length, area), number of grains per spike and TGW for the *TaGS3* mutants were collected using a MARVIN seed analyzer (GTA Sensorik GmbH, Germany), as described previously (Wang *et al*. 2019). The seeds from the three tallest spikes of each plant were analyzed, and the mean values per plant were used for statistical analyses.

### Statistical analysis of data

The distribution phenotypic data was visualized using the boxplot functions from ggplot2 R package (version 3.4.4). The outliers in data were removed using the Rosner’s test for outliers implemented in R package EnvStats (version 2.3.1). One way ANOVA was applied to compare the significance of inter-group differences. The Student’s *t*-test was applied to assess the significance of difference between the reference groups of edited lines (the group with the lowest number of edited *TaGS3* copies in a population) and groups of lines with higher number of the edited *TaGS3* gene loci.

## Data Availability

CRISPR-Cas9 gene editing constructs and wheat lines are available upon request.

## Conflict of Interest

All the authors declare no conflict of interest.

## Authors contribution

W.W. designed/conducted gene editing experiments and developed populations, collected and analyzed phenotypic data, Q.P. analyzed gene editing events using next generation sequencing (NGS); B.T. conducted plant transformation experiments; D.D. collected phenotypic data; G.B. contributed to validation of transgenic constructs by Sanger sequencing; A.A. designed experiments for NGS analysis of editing events and performed NGS; H.T. performed biolistic transformation of wheat embryos with the gene editing constructs; E.A. conceived idea, designed gene editing experiments, coordinated project, analyzed data and wrote the manuscript.

## Acknowledgements

This project was supported by the National Research Initiative Competitive Grants 2021-67013-34174, 2020-67013-30906 and 2022-68013-36439 (WheatCAP) from the USDA National Institute of Food and Agriculture and the Bill and Melinda Gates Foundation grant BMGF:01511000146.

## Notes

### Competing Interest Statement

The authors have declared no competing interest.

## References

Ashikari, M., H. Sakakibara, S. Lin, T. Yamamoto, T. Takashi et al., 2005 Plant science: Cytokinin oxidase regulates rice grain production. Science (80-.). 309: 741–745.

Avni, R., M. Nave, O. Barad, K. Baruch, S. O. Twardziok et al., 2017 Wild emmer genome architecture and diversity elucidate wheat evolution and domestication. Science 97: 93–97.

Benjamini, Y., and Y. Hochberg, 1995 Controlling the False Discovery Rate: A Practical and Powerful Approach to Multiple Testing. J. R. Stat. Soc. Ser. B, Stat. Methodol. 57: 289–300.

Chen, H., N. Patterson, and D. Reich, 2010 Population differentiation as a test for selective sweeps. Genome Res. 20: 393–402.

Edgar, R. C., 2004 MUSCLE: multiple sequence alignment with high accuracy and high throughput. Nucleic Acids Res. 32: 1792–7.

Fan, C., Y. Xing, H. Mao, T. Lu, B. Han et al., 2006 GS3, a major QTL for grain length and weight and minor QTL for grain width and thickness in rice, encodes a putative transmembrane protein. Theor. Appl. Genet. 112: 1164–1171.

Geng, M., Q. Qian, G. Zhang, D. Zeng, R. Xu et al., 2017 Natural Variation in the Promoter of GSE5 Contributes to Grain Size Diversity in Rice. Mol. Plant 10: 685–694.

He, F., R. Pasam, F. Shi, S. Kant, G. Keeble-Gagnere et al., 2019 Exome sequencing highlights the role of wild relative introgression in shaping the adaptive landscape of the wheat genome. Nat. Genet. 51: 896–904.

Hu, J., Y. Wang, Y. Fang, L. Zeng, J. Xu et al., 2015 A Rare Allele of GS2 Enhances Grain Size and Grain Yield in Rice. Mol. Plant 8: 1455–65.

Huang, X., Q. Qian, Z. Liu, H. Sun, S. He et al., 2009 Natural variation at the DEP1 locus enhances grain yield in rice. Nat. Genet. 41: 494–497.

Li, Y., C. Fan, Y. Xing, Y. Jiang, L. Luo et al., 2011 Natural variation in GS5 plays an important role in regulating grain size and yield in rice. Nat. Genet. 43: 1266–9.

Li, N., and Y. Li, 2016 Signaling pathways of seed size control in plants. Curr. Opin. Plant Biol. 33: 23–32.

Li, W., and B. Yang, 2017 Translational genomics of grain size regulation in wheat. Theor. Appl. Genet. 130: 1765–1771.

Liu, L., Y. Zhou, F. Mao, Y. Gu, Z. Tang et al., 2022 Fine-Tuning of the Grain Size by Alternative Splicing of GS3 in Rice. Rice 15:.

Mao, H., S. Sun, J. Yao, C. Wang, S. Yu et al., 2010 Linking differential domain functions of the GS3 protein to natural variation of grain size in rice. Proc. Natl. Acad. Sci. U. S. A. 107: 19579–19584.

Ren, X., L. Zhi, L. Liu, D. Meng, Q. Su et al., 2021 Alternative splicing of tags3 differentially regulates grain weight and size in bread wheat. Int. J. Mol. Sci. 22: 1–20.

Simmonds, J., P. Scott, J. Brinton, T. C. Mestre, M. Bush et al., 2016 A splice acceptor site mutation in TaGW2-A1 increases thousand grain weight in tetraploid and hexaploid wheat through wider and longer grains. Theor. Appl. Genet.

Song, X.-J., W. Huang, M. Shi, M.-Z. Zhu, and H.-X. Lin, 2007 A QTL for rice grain width and weight encodes a previously unknown RING-type E3 ubiquitin ligase. Nat. Genet. 39: 623–30.

Sun, S., L. Wang, H. Mao, L. Shao, X. Li et al., 2018 A G-protein pathway determines grain size in rice. Nat. Commun. 9:.

Takano-Kai, N., J. Hui, T. Kubo, M. Sweeney, T. Matsumoto et al., 2009 Evolutionary history of GS3, a gene conferring grain length in rice. Genetics 182: 1323–1334.

The International Wheat Genome Sequencing Consortium (IWGSC), 2018 Shifting the limits in wheat research and breeding using a fully annotated reference genome. Science 361: eaar7191.

Thung, L., Y. Trusov, D. Chakravorty, and J. R. Botella, 2012 Gγ1+Gγ2+Gγ3=Gβ: The search for heterotrimeric G-protein γ subunits in Arabidopsis is over. J. Plant Physiol. 169: 542–545.

Urano, D., and A. M. Jones, 2014 Heterotrimeric G Protein–Coupled Signaling in Plants. Annu. Rev. Plant Biol. 65: 365–384.

Wang, S., S. Li, Q. Liu, K. Wu, J. Zhang et al., 2015 The OsSPL16-GW7 regulatory module determines grain shape and simultaneously improves rice yield and grain quality. Nat. Genet. 47: 949–954.

Wang, W., Q. Pan, F. He, A. Akhunova, S. Chao et al., 2018a Transgenerational CRISPR-Cas9 Activity Facilitates Multiplex Gene Editing in Allopolyploid Wheat. Cris. J. 1: 65–74.

Wang, W., Q. Pan, B. Tian, F. He, Y. Chen et al., 2019 Gene editing of the wheat homologs of TONNEAU1–recruiting motif encoding gene affects grain shape and weight in wheat. Plant J. 100: 251–264.

Wang, W., J. Simmonds, Q. Pan, D. Davidson, E. Akhunov et al., 2018b Gene editing and mutagenesis reveal inter-cultivar differences and additivity in the contribution of TaGW2 homoeologues to grain size and weight in wheat. Theor. Appl. Genet. 131: 2463–2475.

Wang, W., B. Tian, Q. Pan, Y. Chen, F. He et al., 2021 Expanding the range of editable targets in the wheat genome using the variants of the Cas12a and Cas9 nucleases. Plant Biotechnol. J. 19: 2428–2441.

Wang, S., K. Wu, Q. Yuan, X. Liu, Z. Liu et al., 2012 Control of grain size, shape and quality by OsSPL16 in rice. Nat. Genet. 44: 950–4.

Yang, J., Y. Zhou, Y. Zhang, W. Hu, Q. Wu et al., 2019 Cloning, characterization of TaGS3 and identification of allelic variation associated with kernel traits in wheat (Triticum aestivum L.). BMC Genet. 20: 1–11.

Zhang, W., H. Li, L. Zhi, Q. Su, J. Liu et al., 2020 Functional markers developed from TaGS3, a negative regulator of grain weight and size, for marker-assisted selection in wheat. Crop J. 8: 943–952.

Zhang, L., Y.-L. Zhao, L.-F. Gao, G.-Y. Zhao, R.-H. Zhou et al., 2012 TaCKX6-D1, the ortholog of rice OsCKX2, is associated with grain weight in hexaploid wheat. New Phytol. 195: 574–84.

