## Supplementary Tables for "Non-additive dosage-dependent effects of *TaGS3* gene editing on grain size and weight in wheat"

**Supplementary Materials**

**Supplementary Table 1.** List of oligos used in the project.

| Oligo name | Sequence | Comments |
| --- | --- | --- |
| GS3T6-9checkF | TCCATTGATGACGCTCTCTG | PCR primer |
| GS3T6-9checkR | TCGAGGAAGCTGATCTGG | PCR primer |
| TaGS3T6F | cttGGCCGGCAATGGCGGCGCCC | gRNA oligo |
| TaGS3T6R | aaacGGCCGGCAATGGCGGCGCC | gRNA oligo |
| TaGS3T7F | cttgAAGTCCCCGCTCGACCCCTG | gRNA oligo |
| TaGS3T7R | aaacCAGGGGTCGAGCGGGGACTT | gRNA oligo |
| TaGS3T8F | cttGCGGCCGCAGGGGTCGAGCG | gRNA oligo |
| TaGS3T8R | aaacCGCTCGACCCCTGCGGCCG | gRNA oligo |
| TaGS3T9F | cttGCAGCCGGTGGCGGCCGCAG | gRNA oligo |
| TaGS3T9R | aaacCTGCGGCCGCCACCGGCTG | gRNA oligo |

Supplementary Table 2. Phenotypic data collected for BC1F2 population derived from 4906-1.

| **ID** | **Dosage** | **genotype** | **Seed num.** | **TGW** | **GA** | **GW** | **GL** |
| --- | --- | --- | --- | --- | --- | --- | --- |
| YLD12BC1F1(1)-1-13 | 1 | aaBbdd | 47 | 38.2269504 | 15.1171756 | 3.38332208 | 5.94886265 |
| YLD12BC1F1(1)-1-26 | 1 | aabbDd | 38.3333333 | 49.7391304 | 17.0717504 | 3.77936901 | 6.00332934 |
| YLD12BC1F1(1)-1-33 | 1 | aabbDd | 37.3333333 | 43.3035714 | 16.4175683 | 3.62605928 | 6.01424243 |
| YLD12BC1F1(1)-1-36 | 1 | Aabbdd | 55.3333333 | 39.5783133 | 15.4900036 | 3.51980719 | 5.79632624 |
| YLD12BC1F1(1)-1-4 | 1 | Aabbdd | 40.3333333 | 43.8842975 | 16.9857408 | 3.72775135 | 6.26741489 |
| YLD12BC1F1(1)-1-5 | 1 | aabbDd | 51.3333333 | 42.1428571 | 16.3768465 | 3.58554885 | 6.05582806 |
| YLD12BC1F1(1)-1-73 | 1 | aabbDd | 42.3333333 | 44.8818898 | 17.2609883 | 3.74346801 | 6.37530909 |
| YLD12BC1F1(1)-1-87 | 1 | Aabbdd | 39.6666667 | 35.8823529 | 15.03598 | 3.41446798 | 6.01655367 |
| YLD12BC1F1(1)-1-89 | 1 | aabbDd | 36 | 46.8518519 | 17.5837112 | 3.7961149 | 6.39122638 |
| YLD12BC1F1(1)-5-127 | 1 | aabbDd | 35 | 40.1904762 | 15.7583913 | 3.50624191 | 6.03776283 |
| YLD12BC1F1(1)-5-133 | 1 | aaBbdd | 46 | 39.4927536 | 16.3952218 | 3.5028609 | 6.54993009 |
| YLD12BC1F1(1)-5-141 | 1 | Aabbdd | 49 | 34.6258503 | 14.5095042 | 3.30574417 | 5.90303544 |
| YLD12BC1F1(1)-5-151 | 1 | aabbDd | 42.6666667 | 37.1875 | 15.2195652 | 3.38946044 | 6.14994197 |
| YLD12BC1F1(1)-5-18 | 1 | aabbDd | 48 | 30.4166667 | 13.1675575 | 3.07816262 | 5.73487682 |
| YLD12BC1F1(1)-5-29 | 1 | aaBbdd | 39.3333333 | 37.6271186 | 14.7180529 | 3.30530003 | 6.02556847 |
| YLD12BC1F1(1)-5-50 | 1 | aaBbdd | 45.6666667 | 35.0364964 | 14.8356699 | 3.31039802 | 6.16633969 |
| YLD12BC1F1(1)-5-56 | 1 | aaBbdd | 29.3333333 | 47.0454545 | 17.4703728 | 3.73737783 | 6.29300726 |
| YLD12BC1F1(1)-5-5 | 1 | aabbDd | 55.6666667 | 38.9221557 | 15.5126565 | 3.45099751 | 5.99242911 |
| YLD12BC1F1(1)-5-79 | 1 | Aabbdd | 51.6666667 | 33.4193548 | 14.1882267 | 3.23529264 | 5.93685891 |
| YLD12BC1F1(1)-5-86 | 1 | aaBbdd | 47.6666667 | 42.3776224 | 16.273564 | 3.59480273 | 6.00829039 |
| YLD12BC1F1(1)-5-89 | 1 | aabbDd | 45.3333333 | 38.0882353 | 15.2885299 | 3.44674459 | 5.92645828 |
| YLD12BC1F1(1)-5-8 | 1 | aabbDd | 43.3333333 | 36.6153846 | 14.5897511 | 3.40412503 | 5.73673935 |
| YLD12BC1F1(1)-1-14 | 2 | aaBbDd | 28.6666667 | 43.4883721 | 16.0858064 | 3.69751595 | 5.71166313 |
| YLD12BC1F1(1)-1-15 | 2 | aaBbDd | 40.6666667 | 46.557377 | 17.1783064 | 3.74520245 | 6.07409419 |
| YLD12BC1F1(1)-1-18 | 2 | AabbDd | 38.6666667 | 47.6724138 | 17.2805093 | 3.79220176 | 6.07307869 |
| YLD12BC1F1(1)-1-27 | 2 | aaBbDd | 34.3333333 | 50.0970874 | 17.3901684 | 3.75977355 | 6.22351389 |
| YLD12BC1F1(1)-1-2 | 2 | aabbDD | 39 | 47.4358974 | 17.9907902 | 3.82253527 | 6.45963853 |
| YLD12BC1F1(1)-1-32 | 2 | aaBBdd | 37 | 45.4954955 | 17.4854589 | 3.71662957 | 6.34734451 |
| YLD12BC1F1(1)-1-35 | 2 | AabbDd | 39 | 44.3589744 | 16.6946042 | 3.53980099 | 6.29102632 |
| YLD12BC1F1(1)-1-40 | 2 | AaBbdd | 48.6666667 | 35.7534247 | 14.4956562 | 3.30492882 | 5.80394873 |
| YLD12BC1F1(1)-1-41 | 2 | AabbDd | 42.3333333 | 48.8976378 | 17.7822253 | 3.74650697 | 6.33405351 |
| YLD12BC1F1(1)-1-43 | 2 | AaBbdd | 36.3333333 | 45.5963303 | 16.8124408 | 3.73860564 | 5.96876985 |
| YLD12BC1F1(1)-1-45 | 2 | AabbDd | 38 | 46.6666667 | 17.152427 | 3.70529722 | 6.13125061 |
| YLD12BC1F1(1)-1-50 | 2 | aaBbDd | 45.3333333 | 42.0588235 | 16.2185245 | 3.56500087 | 5.99935424 |
| YLD12BC1F1(1)-1-69 | 2 | AabbDd | 38 | 44.5614035 | 17.2793021 | 3.70396267 | 6.48319443 |
| YLD12BC1F1(1)-1-72 | 2 | AaBbdd | 38.3333333 | 46.9565217 | 17.6949034 | 3.8246832 | 6.34399744 |
| YLD12BC1F1(1)-1-83 | 2 | aaBbDd | 51.3333333 | 38.7012987 | 15.9262731 | 3.43650713 | 6.38478096 |
| YLD12BC1F1(1)-1-92 | 2 | aaBbDd | 39.3333333 | 45.5084746 | 16.3022243 | 3.62545127 | 5.94619931 |
| YLD12BC1F1(1)-1-94 | 2 | aaBbDd | 27 | 41.3580247 | 15.3372774 | 3.51559328 | 5.79193172 |
| YLD12BC1F1(1)-1-95 | 2 | aaBbDd | 37 | 45.6756757 | 16.6398181 | 3.66366702 | 5.99038511 |
| YLD12BC1F1(1)-5-103 | 2 | AaBbdd | 31 | 37.8494624 | 15.3216115 | 3.41879116 | 6.06995669 |
| YLD12BC1F1(1)-5-126 | 2 | AabbDd | 45.3333333 | 37.0588235 | 15.3977894 | 3.41546369 | 6.06504147 |
| YLD12BC1F1(1)-5-129 | 2 | aaBbDd | 43 | 45.3488372 | 16.9158869 | 3.67997975 | 6.14080936 |
| YLD12BC1F1(1)-5-131 | 2 | AabbDd | 52 | 40 | 15.6656189 | 3.46195716 | 6.03276932 |
| YLD12BC1F1(1)-5-140 | 2 | AaBbdd | 47.3333333 | 31.1971831 | 13.919721 | 3.11410234 | 5.97793503 |
| YLD12BC1F1(1)-5-14 | 2 | AAbbdd | 44.3333333 | 36.6917293 | 15.416154 | 3.42164459 | 6.30418843 |
| YLD12BC1F1(1)-5-159 | 2 | aaBbDd | 36 | 32.5925926 | 13.828603 | 3.20707322 | 5.8335604 |
| YLD12BC1F1(1)-5-27 | 2 | aabbDD | 48.6666667 | 40.890411 | 16.2518921 | 3.58672932 | 6.04960043 |
| YLD12BC1F1(1)-5-28 | 2 | aaBbDd | 37.6666667 | 40.7079646 | 16.0701405 | 3.47708194 | 6.15796858 |
| YLD12BC1F1(1)-5-33 | 2 | aaBbDd | 41.6666667 | 39.44 | 15.4344619 | 3.49484276 | 5.86412439 |
| YLD12BC1F1(1)-5-34 | 2 | AabbDd | 36.6666667 | 36.7272727 | 14.753136 | 3.37572615 | 5.85537425 |
| YLD12BC1F1(1)-5-35 | 2 | AAbbdd | 45 | 42.2222222 | 16.1660674 | 3.58078181 | 6.20023586 |
| YLD12BC1F1(1)-5-43 | 2 | aaBbDd | 49.6666667 | 41.0067114 | 16.0473253 | 3.54647777 | 5.96500312 |
| YLD12BC1F1(1)-5-47 | 2 | AaBbdd | 46.3333333 | 37.6978417 | 15.1787668 | 3.38358766 | 5.91714061 |
| YLD12BC1F1(1)-5-48 | 2 | AaBbdd | 39 | 37.3504274 | 15.0692719 | 3.39784214 | 5.93805757 |
| YLD12BC1F1(1)-5-49 | 2 | AabbDd | 49 | 41.3605442 | 16.9541622 | 3.55702841 | 6.71874562 |
| YLD12BC1F1(1)-5-51 | 2 | AaBbdd | 44.3333333 | 36.3909774 | 15.1752683 | 3.39270918 | 6.14993058 |
| YLD12BC1F1(1)-5-55 | 2 | aaBBdd | 42.6666667 | 42.1875 | 16.2179379 | 3.60206509 | 6.00877453 |
| YLD12BC1F1(1)-5-68 | 2 | AabbDd | 47.3333333 | 33.4507042 | 14.6042363 | 3.27302644 | 6.18964926 |
| YLD12BC1F1(1)-5-70 | 2 | AabbDd | 50 | 43.7333333 | 17.082462 | 3.6575799 | 6.39766887 |
| YLD12BC1F1(1)-5-73 | 2 | AaBbdd | 48.6666667 | 38.1506849 | 15.6894459 | 3.47314416 | 6.22202036 |
| YLD12BC1F1(1)-5-77 | 2 | AabbDd | 48.6666667 | 35.9589041 | 14.6705164 | 3.32138774 | 5.85902553 |
| YLD12BC1F1(1)-5-85 | 2 | AabbDd | 29.3333333 | 41.7045455 | 16.0013395 | 3.53113923 | 6.09526685 |
| YLD12BC1F1(1)-5-87 | 2 | aaBBdd | 41 | 39.9186992 | 15.5809192 | 3.47265107 | 5.94398946 |
| YLD12BC1F1(1)-5-88 | 2 | aaBbDd | 56 | 36.7261905 | 14.9358074 | 3.3635761 | 5.92313954 |
| YLD12BC1F1(1)-5-91 | 2 | aaBbDd | 29 | 32.8735632 | 13.8880351 | 3.11817373 | 5.96437199 |
| YLD12BC1F1(1)-5-96 | 2 | AAbbdd | 36.6666667 | 36.0909091 | 14.5037242 | 3.30396818 | 5.93010212 |
| YLD12BC1F1(1)-1-10 | 3 | AaBbDd | 48 | 45.625 | 16.9808256 | 3.77222527 | 5.91116009 |
| YLD12BC1F1(1)-1-16 | 3 | AABbdd | 60.6666667 | 40 | 15.8096369 | 3.47424345 | 6.01667362 |
| YLD12BC1F1(1)-1-17 | 3 | AaBbDd | 35.3333333 | 32.1698113 | 13.380893 | 3.18659608 | 5.57134377 |
| YLD12BC1F1(1)-1-21 | 3 | AaBbDd | 49.3333333 | 44.5945946 | 16.9264022 | 3.65487129 | 6.17131875 |
| YLD12BC1F1(1)-1-22 | 3 | AaBbDd | 38.6666667 | 47.6724138 | 17.1854244 | 3.75821384 | 6.03198175 |
| YLD12BC1F1(1)-1-25 | 3 | aaBBDd | 52 | 39.0384615 | 15.1300649 | 3.44640721 | 5.81543648 |
| YLD12BC1F1(1)-1-30 | 3 | AaBbDd | 43.6666667 | 45.4961832 | 16.3354559 | 3.63800065 | 5.98075561 |
| YLD12BC1F1(1)-1-31 | 3 | AABbdd | 24.3333333 | 55.890411 | 19.9506758 | 4.07692944 | 6.48858371 |
| YLD12BC1F1(1)-1-34 | 3 | AABbdd | 40.6666667 | 45.2459016 | 16.5558351 | 3.64080451 | 6.02145614 |
| YLD12BC1F1(1)-1-37 | 3 | AaBbDd | 21 | 43.6507937 | 16.0945336 | 3.62980389 | 5.88703986 |
| YLD12BC1F1(1)-1-42 | 3 | AaBbDd | 38.3333333 | 43.4782609 | 16.4068558 | 3.62350734 | 6.01494254 |
| YLD12BC1F1(1)-1-44 | 3 | AaBbDd | 49.3333333 | 45.2027027 | 17.2776525 | 3.68384912 | 6.17800803 |
| YLD12BC1F1(1)-1-52 | 3 | AAbbDd | 24.3333333 | 54.9315068 | 18.9008995 | 3.98806047 | 6.29133296 |
| YLD12BC1F1(1)-1-53 | 3 | AaBbDd | 47.3333333 | 45.6338028 | 17.3607821 | 3.69392713 | 6.20552716 |
| YLD12BC1F1(1)-1-55 | 3 | AABbdd | 40 | 49.5833333 | 17.8020095 | 3.84945565 | 6.13256547 |
| YLD12BC1F1(1)-1-61 | 3 | aaBbDD | 49 | 40.4761905 | 16.2868146 | 3.51585826 | 6.2783002 |
| YLD12BC1F1(1)-1-63 | 3 | aaBBDd | 35 | 43.2380952 | 16.0034137 | 3.60986515 | 5.87299951 |
| YLD12BC1F1(1)-1-65 | 3 | aaBbDD | 38 | 43.9473684 | 16.6004532 | 3.66060433 | 6.03323824 |
| YLD12BC1F1(1)-1-66 | 3 | AaBbDd | 37 | 37.5675676 | 14.5470149 | 3.38164361 | 5.76697777 |
| YLD12BC1F1(1)-1-6 | 3 | AAbbDd | 34.6666667 | 44.9038462 | 16.6089178 | 3.66111634 | 6.03372173 |
| YLD12BC1F1(1)-1-70 | 3 | aaBBDd | 50.3333333 | 41.0596026 | 16.4864119 | 3.56796853 | 6.37279322 |
| YLD12BC1F1(1)-1-71 | 3 | AabbDD | 37.3333333 | 46.3392857 | 17.6205275 | 3.79697813 | 6.36727348 |
| YLD12BC1F1(1)-1-77 | 3 | AAbbDd | 46 | 41.4492754 | 17.061797 | 3.67370932 | 6.35141808 |
| YLD12BC1F1(1)-1-79 | 3 | AaBbDd | 36 | 44.3518519 | 16.895781 | 3.65727792 | 6.43084176 |
| YLD12BC1F1(1)-1-80 | 3 | AaBbDd | 38.3333333 | 36.9565217 | 15.2515371 | 3.37403928 | 6.24004257 |
| YLD12BC1F1(1)-1-81 | 3 | AAbbDd | 31.3333333 | 44.0425532 | 16.769228 | 3.70340797 | 6.20624406 |
| YLD12BC1F1(1)-1-82 | 3 | aaBBDd | 33.3333333 | 46.3 | 17.6651176 | 3.77456393 | 6.43285725 |
| YLD12BC1F1(1)-1-91 | 3 | aaBBDd | 34.3333333 | 41.7475728 | 16.3282979 | 3.58578166 | 6.08056606 |
| YLD12BC1F1(1)-1-93 | 3 | AaBbDd | 28.3333333 | 43.7647059 | 16.1789187 | 3.62303625 | 5.8783156 |
| YLD12BC1F1(1)-1-96 | 3 | AaBbDd | 32.6666667 | 43.6734694 | 16.738878 | 3.67411025 | 6.05678067 |
| YLD12BC1F1(1)-5-100 | 3 | AaBbDd | 41 | 38.2113821 | 15.1925528 | 3.39126706 | 5.93686211 |
| YLD12BC1F1(1)-5-101 | 3 | AaBbDd | 34.3333333 | 34.5631068 | 14.184484 | 3.30778874 | 5.75016806 |
| YLD12BC1F1(1)-5-106 | 3 | aaBBDd | 48.3333333 | 36.9655172 | 14.9942548 | 3.27286627 | 6.11946763 |
| YLD12BC1F1(1)-5-107 | 3 | AaBBdd | 42 | 43.8888889 | 17.3523916 | 3.70570264 | 6.48516469 |
| YLD12BC1F1(1)-5-109 | 3 | AAbbDd | 45.3333333 | 36.9117647 | 15.0144227 | 3.32930839 | 5.99299715 |
| YLD12BC1F1(1)-5-10 | 3 | AABbdd | 47 | 44.0425532 | 16.6530686 | 3.58989686 | 6.13546478 |
| YLD12BC1F1(1)-5-116 | 3 | aaBbDD | 45.3333333 | 34.8529412 | 14.7328465 | 3.31178346 | 5.98953468 |
| YLD12BC1F1(1)-5-117 | 3 | aaBbDD | 44.6666667 | 37.761194 | 15.0596049 | 3.40950526 | 5.8453934 |
| YLD12BC1F1(1)-5-124 | 3 | AaBbDd | 34 | 40.5882353 | 16.0471112 | 3.46062484 | 6.21999762 |
| YLD12BC1F1(1)-5-125 | 3 | AaBbDd | 45 | 37.4814815 | 15.6331025 | 3.41480605 | 6.07952528 |
| YLD12BC1F1(1)-5-132 | 3 | AaBbDd | 48 | 38.0555556 | 16.0357643 | 3.46737803 | 6.58218674 |
| YLD12BC1F1(1)-5-134 | 3 | AAbbDd | 46.6666667 | 38.8571429 | 15.5015759 | 3.36637466 | 6.19314021 |
| YLD12BC1F1(1)-5-136 | 3 | AaBBdd | 51.3333333 | 40 | 16.2670487 | 3.51025288 | 6.40008514 |
| YLD12BC1F1(1)-5-139 | 3 | AaBbDd | 45 | 36.8148148 | 15.0768321 | 3.36634421 | 5.98199765 |
| YLD12BC1F1(1)-5-13 | 3 | AaBBdd | 57.6666667 | 39.7687861 | 15.7651309 | 3.48488331 | 6.02820084 |
| YLD12BC1F1(1)-5-143 | 3 | AaBbDd | 49 | 39.047619 | 15.8755701 | 3.42167662 | 6.23551701 |
| YLD12BC1F1(1)-5-149 | 3 | AaBbDd | 58.3333333 | 35.4285714 | 14.7477729 | 3.32372535 | 5.88710399 |
| YLD12BC1F1(1)-5-150 | 3 | AaBBdd | 49.6666667 | 38.0536913 | 15.2603102 | 3.44561868 | 5.93359747 |
| YLD12BC1F1(1)-5-160 | 3 | AAbbDd | 42.6666667 | 33.4375 | 14.2004048 | 3.22731746 | 5.88529614 |
| YLD12BC1F1(1)-5-16 | 3 | aaBbDD | 46.3333333 | 41.7985612 | 16.5358258 | 3.59176913 | 6.1118059 |
| YLD12BC1F1(1)-5-17 | 3 | aaBbDD | 49 | 36.122449 | 14.9230114 | 3.34621468 | 5.97498023 |
| YLD12BC1F1(1)-5-20 | 3 | AaBbDd | 48 | 38.75 | 15.6760056 | 3.44910929 | 6.04122964 |
| YLD12BC1F1(1)-5-21 | 3 | AaBbDd | 41.6666667 | 36.56 | 14.9035907 | 3.37267993 | 5.86721262 |
| YLD12BC1F1(1)-5-24 | 3 | AABbdd | 41.6666667 | 39.76 | 15.6562456 | 3.5295533 | 5.89736254 |
| YLD12BC1F1(1)-5-26 | 3 | aaBBDd | 39.3333333 | 41.779661 | 16.1860492 | 3.54901839 | 6.12079937 |
| YLD12BC1F1(1)-5-2 | 3 | AAbbDd | 50.6666667 | 41.0526316 | 15.7600535 | 3.44784873 | 6.13872156 |
| YLD12BC1F1(1)-5-31 | 3 | AaBbDd | 39 | 42.8205128 | 15.8495728 | 3.54708155 | 5.98098126 |
| YLD12BC1F1(1)-5-36 | 3 | AaBbDd | 32 | 41.3541667 | 16.2738868 | 3.55649037 | 6.15628965 |
| YLD12BC1F1(1)-5-37 | 3 | AaBBdd | 45 | 36.7407407 | 14.978283 | 3.36617712 | 5.93053554 |
| YLD12BC1F1(1)-5-39 | 3 | aaBbDD | 42.3333333 | 39.8425197 | 15.9123829 | 3.49103548 | 6.12089243 |
| YLD12BC1F1(1)-5-46 | 3 | AaBbDd | 24.3333333 | 38.630137 | 15.9168256 | 3.54987436 | 6.06878345 |
| YLD12BC1F1(1)-5-53 | 3 | AaBbDd | 43 | 41.7054264 | 16.5625871 | 3.6091866 | 6.32968754 |
| YLD12BC1F1(1)-5-54 | 3 | aaBbDD | 35.6666667 | 42.8971963 | 16.6458912 | 3.59064373 | 6.21761564 |
| YLD12BC1F1(1)-5-58 | 3 | AabbDD | 42.3333333 | 42.8346457 | 16.5107014 | 3.54720245 | 6.19497214 |
| YLD12BC1F1(1)-5-65 | 3 | AaBbDd | 22 | 33.9393939 | 14.5299566 | 3.28873897 | 5.98352002 |
| YLD12BC1F1(1)-5-69 | 3 | AAbbDd | 41 | 41.7886179 | 16.0283372 | 3.53314847 | 6.03730568 |
| YLD12BC1F1(1)-5-71 | 3 | AabbDD | 34 | 39.9019608 | 16.0641203 | 3.5324638 | 6.27998275 |
| YLD12BC1F1(1)-5-75 | 3 | AabbDD | 53.6666667 | 42.0496894 | 16.743245 | 3.61385192 | 6.36272235 |
| YLD12BC1F1(1)-5-82 | 3 | aaBbDD | 44.3333333 | 37.3684211 | 15.2932779 | 3.38339047 | 6.07190482 |
| YLD12BC1F1(1)-5-83 | 3 | AaBBdd | 44.6666667 | 44.7014925 | 17.4651113 | 3.6786361 | 6.56965328 |
| YLD12BC1F1(1)-5-95 | 3 | aaBbDD | 42 | 39.6825397 | 15.4905106 | 3.48001471 | 5.93608055 |
| YLD12BC1F1(1)-5-97 | 3 | AaBBdd | 58 | 40.4597701 | 16.1971454 | 3.54900069 | 6.21016278 |
| YLD12BC1F1(1)-1-11 | 4 | AABbDd | 46.3333333 | 40.5035971 | 15.8378949 | 3.44572951 | 6.12993261 |
| YLD12BC1F1(1)-1-1 | 4 | AABbDd | 55 | 38.8484848 | 15.887339 | 3.41978616 | 6.23505149 |
| YLD12BC1F1(1)-1-20 | 4 | AaBBDd | 41 | 40.0813008 | 15.79922 | 3.48380905 | 6.13109594 |
| YLD12BC1F1(1)-1-24 | 4 | AaBBDd | 31.3333333 | 38.4042553 | 14.4882442 | 3.37472015 | 5.74747627 |
| YLD12BC1F1(1)-1-28 | 4 | AABbDd | 35.3333333 | 48.2075472 | 16.8031823 | 3.66497493 | 6.12197415 |
| YLD12BC1F1(1)-1-29 | 4 | AABbDd | 55.3333333 | 41.0843373 | 15.7327685 | 3.43804027 | 6.11268309 |
| YLD12BC1F1(1)-1-46 | 4 | AaBbDD | 48.6666667 | 41.8493151 | 16.4778557 | 3.54390737 | 6.19939445 |
| YLD12BC1F1(1)-1-48 | 4 | AaBBDd | 43.6666667 | 39.389313 | 15.5874923 | 3.50196005 | 5.90134057 |
| YLD12BC1F1(1)-1-49 | 4 | AABbDd | 53 | 39.6226415 | 15.7050194 | 3.50959692 | 5.89135143 |
| YLD12BC1F1(1)-1-68 | 4 | AABBdd | 36.6666667 | 45.1818182 | 17.5581204 | 3.7098765 | 6.65901126 |
| YLD12BC1F1(1)-1-74 | 4 | AaBbDD | 37.3333333 | 44.7321429 | 16.9115777 | 3.71508362 | 6.29016566 |
| YLD12BC1F1(1)-1-75 | 4 | AaBBDd | 28.6666667 | 47.6744186 | 18.3044879 | 3.70761635 | 6.77301858 |
| YLD12BC1F1(1)-1-78 | 4 | AaBbDD | 37.6666667 | 46.460177 | 17.4579909 | 3.75533359 | 6.44741079 |
| YLD12BC1F1(1)-1-88 | 4 | AaBbDD | 40 | 38.5833333 | 16.0722458 | 3.50528902 | 6.39800866 |
| YLD12BC1F1(1)-1-9 | 4 | AAbbDD | 42 | 45.3174603 | 17.3536758 | 3.6991425 | 6.42207611 |
| YLD12BC1F1(1)-5-104 | 4 | AaBbDD | 50.3333333 | 36.6225166 | 14.8160714 | 3.38465482 | 5.87852794 |
| YLD12BC1F1(1)-5-110 | 4 | AABbDd | 30.3333333 | 37.6923077 | 15.4830096 | 3.36416312 | 6.1870734 |
| YLD12BC1F1(1)-5-112 | 4 | AABBdd | 45 | 40.3703704 | 16.1689306 | 3.53462107 | 6.26391828 |
| YLD12BC1F1(1)-5-113 | 4 | AABBdd | 43.6666667 | 41.0687023 | 16.2494744 | 3.54636723 | 6.2412893 |
| YLD12BC1F1(1)-5-115 | 4 | AaBBDd | 42.3333333 | 35.1181102 | 14.7540389 | 3.27284876 | 6.08541741 |
| YLD12BC1F1(1)-5-118 | 4 | AABbDd | 45.3333333 | 39.3382353 | 15.5642895 | 3.53612213 | 5.92697948 |
| YLD12BC1F1(1)-5-119 | 4 | AaBBDd | 44.3333333 | 36.9172932 | 15.516726 | 3.38472279 | 6.29351366 |
| YLD12BC1F1(1)-5-12 | 4 | AaBBDd | 44 | 40.7575758 | 15.857579 | 3.49727957 | 6.05421853 |
| YLD12BC1F1(1)-5-135 | 4 | AABbDd | 56.6666667 | 36.9411765 | 15.346828 | 3.38103559 | 6.09512185 |
| YLD12BC1F1(1)-5-138 | 4 | AABBdd | 55.3333333 | 39.0963855 | 16.0296858 | 3.45645378 | 6.27132238 |
| YLD12BC1F1(1)-5-142 | 4 | AaBBDd | 52.6666667 | 32.721519 | 14.2434115 | 3.22222974 | 5.90203726 |
| YLD12BC1F1(1)-5-144 | 4 | aaBBDD | 51.3333333 | 33.7662338 | 14.5147215 | 3.23913188 | 5.98648688 |
| YLD12BC1F1(1)-5-145 | 4 | AABbDd | 40.6666667 | 38.1147541 | 15.2923245 | 3.3768946 | 6.04731657 |
| YLD12BC1F1(1)-5-147 | 4 | AABBdd | 47 | 39.787234 | 16.0113228 | 3.46558072 | 6.19284698 |
| YLD12BC1F1(1)-5-148 | 4 | AABbDd | 45 | 34.5925926 | 14.5188971 | 3.26163035 | 5.91979327 |
| YLD12BC1F1(1)-5-158 | 4 | AABbDd | 41 | 36.4227642 | 15.1998612 | 3.31192045 | 6.11810913 |
| YLD12BC1F1(1)-5-15 | 4 | AaBBDd | 55.6666667 | 39.1017964 | 15.7358277 | 3.45685497 | 6.0294237 |
| YLD12BC1F1(1)-5-1 | 4 | AaBBDd | 41 | 36.8292683 | 14.4756754 | 3.3294707 | 5.802922 |
| YLD12BC1F1(1)-5-22 | 4 | AaBBDd | 45.3333333 | 36.7647059 | 14.9793249 | 3.33179288 | 6.04124724 |
| YLD12BC1F1(1)-5-25 | 4 | AaBBDd | 37.6666667 | 40.1769912 | 15.4781998 | 3.48919628 | 5.9162474 |
| YLD12BC1F1(1)-5-41 | 4 | AABbDd | 48 | 38.4722222 | 15.5705063 | 3.3942687 | 6.09385097 |
| YLD12BC1F1(1)-5-42 | 4 | AaBbDD | 34.6666667 | 45.5769231 | 17.3327282 | 3.69814944 | 6.23902329 |
| YLD12BC1F1(1)-5-45 | 4 | AaBbDD | 33.3333333 | 37.7 | 14.8515606 | 3.43189195 | 5.76919718 |
| YLD12BC1F1(1)-5-4 | 4 | AaBbDD | 41.6666667 | 39.12 | 15.6336646 | 3.51606783 | 5.92609042 |
| YLD12BC1F1(1)-5-52 | 4 | AaBBDd | 31.6666667 | 39.6842105 | 16.1578176 | 3.50439077 | 6.60514218 |
| YLD12BC1F1(1)-5-59 | 4 | AaBbDD | 27.6666667 | 34.8192771 | 14.0924943 | 3.27810121 | 5.81282459 |
| YLD12BC1F1(1)-5-67 | 4 | AABbDd | 46 | 40.8695652 | 16.7828686 | 3.52238277 | 6.72460188 |
| YLD12BC1F1(1)-5-6 | 4 | AaBBDd | 44.3333333 | 44.2105263 | 16.7080431 | 3.63943929 | 6.12330552 |
| YLD12BC1F1(1)-5-72 | 4 | AABbDd | 44.6666667 | 43.7313433 | 17.2264927 | 3.63958457 | 6.46479567 |
| YLD12BC1F1(1)-5-84 | 4 | aaBBDD | 49.6666667 | 36.2416107 | 15.1005542 | 3.43247363 | 6.11263708 |
| YLD12BC1F1(1)-5-92 | 4 | AABbDd | 44.3333333 | 37.518797 | 14.7172877 | 3.36020658 | 5.85188239 |
| YLD12BC1F1(1)-5-93 | 4 | AaBbDD | 35.3333333 | 43.0188679 | 16.4715115 | 3.63321591 | 6.01317923 |
| YLD12BC1F1(1)-5-99 | 4 | AAbbDD | 55 | 43.4545455 | 17.0469569 | 3.61807209 | 6.36435069 |
| YLD12BC1F1(1)-1-19 | 5 | AABbDD | 54 | 38.1481481 | 15.4165717 | 3.45041769 | 5.92035121 |
| YLD12BC1F1(1)-1-51 | 5 | AABbDD | 28.3333333 | 50.2352941 | 17.8635111 | 3.910096 | 6.06593422 |
| YLD12BC1F1(1)-1-84 | 5 | AABBDd | 32.6666667 | 43.9795918 | 16.8675715 | 3.64287513 | 6.30924422 |
| YLD12BC1F1(1)-5-105 | 5 | AABbDD | 46.6666667 | 38.7857143 | 15.7571286 | 3.44133593 | 6.15638411 |
| YLD12BC1F1(1)-5-108 | 5 | AABBDd | 43.3333333 | 36.3076923 | 15.0937043 | 3.43040553 | 5.90393868 |
| YLD12BC1F1(1)-5-111 | 5 | AaBBDD | 55.6666667 | 37.005988 | 15.3324939 | 3.38782969 | 6.23847301 |
| YLD12BC1F1(1)-5-114 | 5 | AABBDd | 34 | 43.627451 | 16.7042986 | 3.6491954 | 6.15462086 |
| YLD12BC1F1(1)-5-11 | 5 | AABBDd | 43.6666667 | 41.1450382 | 15.9523476 | 3.45928112 | 6.19882019 |
| YLD12BC1F1(1)-5-122 | 5 | AaBBDD | 50.6666667 | 39.7368421 | 16.0612537 | 3.4749641 | 6.08543272 |
| YLD12BC1F1(1)-5-128 | 5 | AABBDd | 37 | 46.9369369 | 17.5227929 | 3.60935052 | 6.51370075 |
| YLD12BC1F1(1)-5-130 | 5 | AABbDD | 50.3333333 | 45.1655629 | 17.1536432 | 3.68504387 | 6.20776908 |
| YLD12BC1F1(1)-5-137 | 5 | AABBDd | 27 | 45.4320988 | 17.5650667 | 3.73934552 | 6.22622939 |
| YLD12BC1F1(1)-5-152 | 5 | AABBDd | 51.6666667 | 38.9677419 | 16.0337563 | 3.46324297 | 6.37749462 |
| YLD12BC1F1(1)-5-156 | 5 | AaBBDD | 44.3333333 | 34.6616541 | 14.9706099 | 3.31968334 | 6.30603399 |
| YLD12BC1F1(1)-5-157 | 5 | AABBDd | 50 | 39.2666667 | 15.794417 | 3.49413373 | 6.05177267 |
| YLD12BC1F1(1)-5-19 | 5 | AABbDD | 33 | 37.6767677 | 15.0726735 | 3.40589373 | 5.91394909 |
| YLD12BC1F1(1)-5-30 | 5 | AABBDd | 47.3333333 | 39.6478873 | 15.3940274 | 3.41260681 | 5.99166281 |
| YLD12BC1F1(1)-5-32 | 5 | AABbDD | 42 | 33.4920635 | 13.9724198 | 3.1851778 | 5.86044796 |
| YLD12BC1F1(1)-5-38 | 5 | AABBDd | 40.6666667 | 39.4262295 | 15.5704029 | 3.46676184 | 5.98092805 |
| YLD12BC1F1(1)-5-3 | 5 | AaBBDD | 43 | 39.4573643 | 16.2614808 | 3.49087233 | 6.37358208 |
| YLD12BC1F1(1)-5-44 | 5 | AaBBDD | 23.6666667 | 41.6901408 | 17.5007491 | 3.61650369 | 6.76179573 |
| YLD12BC1F1(1)-5-62 | 5 | AaBBDD | 29 | 36.8965517 | 15.533178 | 3.45580903 | 6.28761859 |
| YLD12BC1F1(1)-5-80 | 5 | AABbDD | 43.3333333 | 39.6923077 | 16.3812468 | 3.5047499 | 6.53247272 |
| YLD12BC1F1(1)-5-81 | 5 | AABbDD | 49 | 37.8911565 | 15.6673488 | 3.43495853 | 6.32994463 |
| YLD12BC1F1(1)-5-90 | 5 | AaBBDD | 45.6666667 | 41.6788321 | 17.0164579 | 3.573764 | 6.73666147 |
| YLD12BC1F1(1)-5-94 | 5 | AABBDd | 28 | 37.8571429 | 15.373714 | 3.37581737 | 6.033803 |
| YLD12BC1F1(1)-5-98 | 5 | AaBBDD | 44 | 35.9848485 | 15.3404226 | 3.43251717 | 6.23829199 |

**Supplementary Table 3.** Phenotypic data collected for BC1F3 population derived from 4906-1.

| ID | Dosage | genotype | seedNum_sp | TGW | GA | GW | GL |
| --- | --- | --- | --- | --- | --- | --- | --- |
| YLD12BC1F1(1)_5-23-1 | 0 | aabbdd | 43 | 48.99225 | 18.09906 | 3.863173 | 6.273047 |
| YLD12BC1F1(1)_5-23-10 | 0 | aabbdd | 50 | 39.6 | 16.02892 | 3.550399 | 6.102424 |
| YLD12BC1F1(1)_5-23-2 | 0 | aabbdd | 47.33333 | 40.42254 | 15.78614 | 3.566155 | 5.992914 |
| YLD12BC1F1(1)_5-23-3 | 0 | aabbdd | 46 | 44.78261 | 17.00786 | 3.74884 | 6.045925 |
| YLD12BC1F1(1)_5-23-4 | 0 | aabbdd | 43.66667 | 38.39695 | 15.53363 | 3.481316 | 6.008232 |
| YLD12BC1F1(1)_5-23-5 | 0 | aabbdd | 46.33333 | 41.79856 | 16.2311 | 3.653864 | 5.945587 |
| YLD12BC1F1(1)_5-23-6 | 0 | aabbdd | 40.66667 | 41.47541 | 16.13054 | 3.60627 | 6.011324 |
| YLD12BC1F1(1)_5-23-7 | 0 | aabbdd | 44.66667 | 42.46269 | 16.45234 | 3.625008 | 6.063108 |
| YLD12BC1F1(1)_5-23-8 | 0 | aabbdd | 42.33333 | 43.46457 | 16.6609 | 3.694324 | 6.039682 |
| YLD12BC1F1(1)_5-23-9 | 0 | aabbdd | 48.33333 | 38.41379 | 15.49146 | 3.508576 | 5.937035 |
| YLD12BC1F1(1)_5-79-1 | 0 | aabbdd | 36.33333 | 40 | 15.92283 | 3.504697 | 6.282413 |
| YLD12BC1F1(1)_5-79-11 | 0 | aabbdd | 57.66667 | 39.9422 | 16.21174 | 3.533228 | 6.141313 |
| YLD12BC1F1(1)_5-79-14 | 0 | aabbdd | 46.66667 | 34.92857 | 14.79849 | 3.370308 | 5.976383 |
| YLD12BC1F1(1)_5-79-16 | 0 | aabbdd | 49 | 35.37415 | 14.73051 | 3.349772 | 5.983192 |
| YLD12BC1F1(1)_5-79-20 | 0 | aabbdd | 33.33333 | 41.8 | 16.23906 | 3.528226 | 6.247488 |
| YLD12BC1F1(1)_5-79-23 | 0 | aabbdd | 51.33333 | 39.61039 | 15.83539 | 3.483933 | 6.14351 |
| YLD12BC1F1(1)_5-79-3 | 0 | aabbdd | 47.66667 | 38.67133 | 15.75539 | 3.518915 | 6.09456 |
| YLD12BC1F1(1)_5-79-4 | 0 | aabbdd | 44.33333 | 35.26316 | 14.64348 | 3.363168 | 5.91051 |
| YLD12BC1F1(1)_5-96-16 | 0 | aabbdd | 44 | 41.74242 | 16.631 | 3.590623 | 6.370318 |
| YLD12BC1F1(1)_5-96-17 | 0 | aabbdd | 53.33333 | 43.5625 | 17.01779 | 3.684161 | 6.17241 |
| YLD12BC1F1(1)_5-96-2 | 0 | aabbdd | 48 | 41.04167 | 16.28391 | 3.56573 | 6.162188 |
| YLD12BC1F1(1)_5-96-21 | 0 | aabbdd | 43.66667 | 46.94656 | 17.71926 | 3.740541 | 6.3835 |
| YLD12BC1F1(1)_5-96-22 | 0 | aabbdd | 52.33333 | 42.61146 | 16.56434 | 3.626801 | 6.120963 |
| YLD12BC1F1(1)_5-96-27 | 0 | aabbdd | 50 | 46.93333 | 17.63962 | 3.734867 | 6.354185 |
| YLD12BC1F1(1)_5-96-39 | 0 | aabbdd | 57.33333 | 48.13953 | 18.18757 | 3.799224 | 6.468137 |
| YLD12BC1F1(1)_5-96-7 | 0 | aabbdd | 53.33333 | 42.4375 | 16.62529 | 3.640803 | 6.16844 |
| YLD12BC1F1(1)_5-96-8 | 0 | aabbdd | 42 | 40.47619 | 15.88093 | 3.519954 | 6.132414 |
| YLD12BC1F1(1)_5-79-10 | 1 | Aabbdd | 51.33333 | 40 | 16.52056 | 3.585666 | 6.18472 |
| YLD12BC1F1(1)_5-79-13 | 1 | Aabbdd | 47.33333 | 41.12676 | 16.86295 | 3.5651 | 6.48974 |
| YLD12BC1F1(1)_5-79-15 | 1 | Aabbdd | 46.33333 | 41.94245 | 17.12058 | 3.555858 | 6.491656 |
| YLD12BC1F1(1)_5-79-19 | 1 | Aabbdd | 51.33333 | 35.71429 | 15.34897 | 3.375972 | 6.128085 |
| YLD12BC1F1(1)_5-79-22 | 1 | Aabbdd | 41.33333 | 37.98387 | 15.59736 | 3.436908 | 6.205878 |
| YLD12BC1F1(1)_5-79-25 | 1 | Aabbdd | 39 | 37.69231 | 15.76206 | 3.442519 | 6.292493 |
| YLD12BC1F1(1)_5-79-26 | 1 | Aabbdd | 44.66667 | 42.53731 | 17.23978 | 3.647865 | 6.520415 |
| YLD12BC1F1(1)_5-79-27 | 1 | Aabbdd | 43 | 44.57364 | 17.65635 | 3.712312 | 6.483109 |
| YLD12BC1F1(1)_5-79-28 | 1 | Aabbdd | 41 | 46.66667 | 18.04311 | 3.756334 | 6.515928 |
| YLD12BC1F1(1)_5-79-29 | 1 | Aabbdd | 49.33333 | 40 | 16.31738 | 3.543911 | 6.263834 |
| YLD12BC1F1(1)_5-79-31 | 1 | Aabbdd | 48 | 37.84722 | 15.54279 | 3.424131 | 6.162246 |
| YLD12BC1F1(1)_5-79-32 | 1 | Aabbdd | 52.33333 | 37.32484 | 15.51288 | 3.401352 | 6.147847 |
| YLD12BC1F1(1)_5-79-33 | 1 | Aabbdd | 52.66667 | 37.1519 | 15.86052 | 3.479386 | 6.219485 |
| YLD12BC1F1(1)_5-79-34 | 1 | Aabbdd | 49.33333 | 35.60811 | 15.33178 | 3.39974 | 6.252847 |
| YLD12BC1F1(1)_5-79-35 | 1 | Aabbdd | 39.33333 | 43.64407 | 17.18965 | 3.592002 | 6.446813 |
| YLD12BC1F1(1)_5-79-40 | 1 | Aabbdd | 54.66667 | 39.69512 | 16.34614 | 3.493104 | 6.364688 |
| YLD12BC1F1(1)_5-79-5 | 1 | Aabbdd | 51.33333 | 41.88312 | 16.51887 | 3.561253 | 6.194959 |
| YLD12BC1F1(1)_5-79-7 | 1 | Aabbdd | 53 | 36.66667 | 15.24996 | 3.431138 | 6.021454 |
| YLD12BC1F1(1)_5-79-8 | 1 | Aabbdd | 40 | 37.08333 | 15.47708 | 3.416355 | 6.128001 |
| YLD12BC1F1(1)_5-96-1 | 1 | Aabbdd | 46 | 45.57971 | 17.45348 | 3.749031 | 6.274915 |
| YLD12BC1F1(1)_5-96-10 | 1 | Aabbdd | 49 | 46.80272 | 18.01692 | 3.734339 | 6.413339 |
| YLD12BC1F1(1)_5-96-11 | 1 | Aabbdd | 49.33333 | 45.13514 | 17.61396 | 3.706447 | 6.345819 |
| YLD12BC1F1(1)_5-96-15 | 1 | Aabbdd | 49.66667 | 48.45638 | 18.77072 | 3.84012 | 6.598383 |
| YLD12BC1F1(1)_5-96-25 | 1 | Aabbdd | 49.66667 | 45.97315 | 17.90956 | 3.733287 | 6.46357 |
| YLD12BC1F1(1)_5-96-28 | 1 | Aabbdd | 43 | 52.63566 | 19.31003 | 3.932336 | 6.647435 |
| YLD12BC1F1(1)_5-96-29 | 1 | Aabbdd | 43 | 43.10078 | 16.99296 | 3.657076 | 6.386646 |
| YLD12BC1F1(1)_5-96-3 | 1 | Aabbdd | 45 | 45.77778 | 17.37829 | 3.722387 | 6.29735 |
| YLD12BC1F1(1)_5-96-32 | 1 | Aabbdd | 58.33333 | 45.31429 | 17.71943 | 3.717304 | 6.437189 |
| YLD12BC1F1(1)_5-96-34 | 1 | Aabbdd | 47 | 50 | 18.74923 | 3.866689 | 6.630767 |
| YLD12BC1F1(1)_5-96-36 | 1 | Aabbdd | 44 | 44.62121 | 17.41722 | 3.649415 | 6.508024 |
| YLD12BC1F1(1)_5-96-4 | 1 | Aabbdd | 46 | 44.42029 | 17.32757 | 3.702316 | 6.308485 |
| YLD12BC1F1(1)_5-96-40 | 1 | Aabbdd | 39.66667 | 47.56303 | 18.04732 | 3.706362 | 6.587954 |
| YLD12BC1F1(1)_5-96-9 | 1 | Aabbdd | 47.33333 | 43.73239 | 17.30204 | 3.687868 | 6.30907 |
| YLD12BC1F1(1)_1-2-1 | 2 | aabbDD | 52.33333 | 44.01274 | 17.1523 | 3.679115 | 6.239526 |
| YLD12BC1F1(1)_1-2-10 | 2 | aabbDD | 51.33333 | 49.35065 | 18.89426 | 3.865969 | 6.755537 |
| YLD12BC1F1(1)_1-2-2 | 2 | aabbDD | 40.33333 | 58.18182 | 20.47202 | 4.057241 | 6.686157 |
| YLD12BC1F1(1)_1-2-3 | 2 | aabbDD | 30.33333 | 57.25275 | 20.2659 | 4.028175 | 6.762449 |
| YLD12BC1F1(1)_1-2-4 | 2 | aabbDD | 39.33333 | 54.0678 | 19.32979 | 3.959732 | 6.56416 |
| YLD12BC1F1(1)_1-2-5 | 2 | aabbDD | 35.33333 | 49.33962 | 18.7916 | 3.907246 | 6.418157 |
| YLD12BC1F1(1)_1-2-6 | 2 | aabbDD | 51 | 46.79739 | 18.08721 | 3.738862 | 6.515716 |
| YLD12BC1F1(1)_1-2-7 | 2 | aabbDD | 35 | 55.04762 | 19.72361 | 3.968489 | 6.730855 |
| YLD12BC1F1(1)_1-2-9 | 2 | aabbDD | 44.66667 | 49.1791 | 18.17392 | 3.772256 | 6.52879 |
| YLD12BC1F1(1)_5-27-1 | 2 | aabbDD | 47.33333 | 46.12676 | 17.90071 | 3.793307 | 6.345217 |
| YLD12BC1F1(1)_5-27-10 | 2 | aabbDD | 52.33333 | 44.4586 | 17.40561 | 3.677189 | 6.444603 |
| YLD12BC1F1(1)_5-27-2 | 2 | aabbDD | 47.33333 | 45.98592 | 17.82816 | 3.752687 | 6.37963 |
| YLD12BC1F1(1)_5-27-3 | 2 | aabbDD | 50 | 42.33333 | 16.94912 | 3.667734 | 6.221622 |
| YLD12BC1F1(1)_5-27-4 | 2 | aabbDD | 50 | 45.4 | 18.03221 | 3.761467 | 6.495513 |
| YLD12BC1F1(1)_5-27-5 | 2 | aabbDD | 51.33333 | 44.35065 | 17.51557 | 3.750965 | 6.330395 |
| YLD12BC1F1(1)_5-27-6 | 2 | aabbDD | 53.66667 | 46.08696 | 17.586 | 3.769088 | 6.367974 |
| YLD12BC1F1(1)_5-27-7 | 2 | aabbDD | 49 | 42.58503 | 17.60482 | 3.631139 | 6.61831 |
| YLD12BC1F1(1)_5-27-8 | 2 | aabbDD | 48 | 46.52778 | 17.75228 | 3.770411 | 6.410874 |
| YLD12BC1F1(1)_5-27-9 | 2 | aabbDD | 44 | 51.66667 | 19.13642 | 3.866625 | 6.721044 |
| YLD12BC1F1(1)_1-32-1 | 2 | aaBBdd | 36.66667 | 55 | 19.32296 | 3.969079 | 6.549551 |
| YLD12BC1F1(1)_1-32-10 | 2 | aaBBdd | 42.33333 | 52.44094 | 19.04172 | 3.896059 | 6.666267 |
| YLD12BC1F1(1)_1-32-2 | 2 | aaBBdd | 31.33333 | 54.25532 | 19.10549 | 4.008892 | 6.433235 |
| YLD12BC1F1(1)_1-32-4 | 2 | aaBBdd | 41.66667 | 46.16 | 17.50976 | 3.716433 | 6.349339 |
| YLD12BC1F1(1)_1-32-6 | 2 | aaBBdd | 35.33333 | 52.45283 | 19.02693 | 3.863349 | 6.62532 |
| YLD12BC1F1(1)_1-32-7 | 2 | aaBBdd | 48 | 42.43056 | 16.81964 | 3.651678 | 6.188878 |
| YLD12BC1F1(1)_1-32-8 | 2 | aaBBdd | 30.33333 | 50 | 18.15056 | 3.843683 | 6.31313 |
| YLD12BC1F1(1)_1-32-9 | 2 | aaBBdd | 47.66667 | 46.64336 | 17.67495 | 3.76747 | 6.361464 |
| YLD12BC1F1(1)_5-55-1 | 2 | aaBBdd | 52 | 45.32051 | 17.58333 | 3.763966 | 6.24079 |
| YLD12BC1F1(1)_5-55-10 | 2 | aaBBdd | 56 | 46.30952 | 17.59407 | 3.719566 | 6.389604 |
| YLD12BC1F1(1)_5-55-2 | 2 | aaBBdd | 46 | 44.71014 | 17.75045 | 3.702318 | 6.496853 |
| YLD12BC1F1(1)_5-55-3 | 2 | aaBBdd | 34.33333 | 49.51456 | 18.58121 | 3.753905 | 6.671317 |
| YLD12BC1F1(1)_5-55-4 | 2 | aaBBdd | 52.33333 | 45.22293 | 17.52373 | 3.74261 | 6.324923 |
| YLD12BC1F1(1)_5-55-5 | 2 | aaBBdd | 40 | 49.5 | 18.99908 | 3.820023 | 6.731164 |
| YLD12BC1F1(1)_5-55-6 | 2 | aaBBdd | 49.66667 | 46.10738 | 18.41639 | 3.759148 | 6.74163 |
| YLD12BC1F1(1)_5-55-7 | 2 | aaBBdd | 43.66667 | 43.43511 | 17.59087 | 3.68596 | 6.552921 |
| YLD12BC1F1(1)_5-55-8 | 2 | aaBBdd | 58 | 46.43678 | 17.62448 | 3.767986 | 6.306045 |
| YLD12BC1F1(1)_5-55-9 | 2 | aaBBdd | 44 | 49.01515 | 18.68478 | 3.888557 | 6.514747 |
| YLD12BC1F1(1)_5-14-1 | 2 | AAbbdd | 42 | 38.49206 | 15.72226 | 3.412426 | 6.265229 |
| YLD12BC1F1(1)_5-14-10 | 2 | AAbbdd | 41.33333 | 45.16129 | 17.19665 | 3.672426 | 6.36429 |
| YLD12BC1F1(1)_5-14-2 | 2 | AAbbdd | 36.33333 | 38.53211 | 15.88242 | 3.466755 | 6.210809 |
| YLD12BC1F1(1)_5-14-3 | 2 | AAbbdd | 46.66667 | 44.35714 | 17.35202 | 3.626438 | 6.460313 |
| YLD12BC1F1(1)_5-14-4 | 2 | AAbbdd | 47 | 37.80142 | 16.0212 | 3.439877 | 6.337953 |
| YLD12BC1F1(1)_5-14-5 | 2 | AAbbdd | 46.33333 | 50.14388 | 18.7975 | 3.841064 | 6.574492 |
| YLD12BC1F1(1)_5-14-6 | 2 | AAbbdd | 54.33333 | 41.34969 | 16.47692 | 3.596389 | 6.221464 |
| YLD12BC1F1(1)_5-14-7 | 2 | AAbbdd | 47.33333 | 41.83099 | 17.44859 | 3.619879 | 6.528632 |
| YLD12BC1F1(1)_5-14-8 | 2 | AAbbdd | 48.33333 | 45.86207 | 17.88285 | 3.762113 | 6.427454 |
| YLD12BC1F1(1)_5-14-9 | 2 | AAbbdd | 43 | 44.49612 | 17.09769 | 3.633316 | 6.39938 |
| YLD12BC1F1(1)_5-35-1 | 2 | AAbbdd | 54 | 43.45679 | 17.14235 | 3.67847 | 6.217166 |
| YLD12BC1F1(1)_5-35-2 | 2 | AAbbdd | 46 | 46.5942 | 17.59353 | 3.702534 | 6.519624 |
| YLD12BC1F1(1)_5-35-3 | 2 | AAbbdd | 40 | 51.91667 | 19.28022 | 3.931069 | 6.653698 |
| YLD12BC1F1(1)_5-35-4 | 2 | AAbbdd | 52.33333 | 45.41401 | 17.7811 | 3.771021 | 6.297275 |
| YLD12BC1F1(1)_5-35-5 | 2 | AAbbdd | 45.66667 | 46.71533 | 18.1041 | 3.785144 | 6.410619 |
| YLD12BC1F1(1)_5-35-6 | 2 | AAbbdd | 51.66667 | 48 | 18.19635 | 3.768173 | 6.485568 |
| YLD12BC1F1(1)_5-35-7 | 2 | AAbbdd | 55 | 44.06061 | 17.3825 | 3.723219 | 6.228315 |
| YLD12BC1F1(1)_5-35-8 | 2 | AAbbdd | 51 | 48.16993 | 18.16729 | 3.7628 | 6.454232 |
| YLD12BC1F1(1)_5-35-9 | 2 | AAbbdd | 44.33333 | 49.09774 | 18.63574 | 3.832632 | 6.613785 |
| YLD12BC1F1(1)_5-79-12 | 2 | AAbbdd | 50.66667 | 36.51316 | 15.78483 | 3.428207 | 6.218843 |
| YLD12BC1F1(1)_5-79-17 | 2 | AAbbdd | 51.66667 | 38 | 15.73595 | 3.443362 | 6.253976 |
| YLD12BC1F1(1)_5-79-18 | 2 | AAbbdd | 60 | 44.77778 | 17.38338 | 3.69379 | 6.372162 |
| YLD12BC1F1(1)_5-79-21 | 2 | AAbbdd | 49.66667 | 39.79866 | 15.94543 | 3.479836 | 6.184431 |
| YLD12BC1F1(1)_5-79-24 | 2 | AAbbdd | 47 | 40.35461 | 16.24874 | 3.486335 | 6.351234 |
| YLD12BC1F1(1)_5-79-30 | 2 | AAbbdd | 48 | 40 | 16.36887 | 3.472048 | 6.446423 |
| YLD12BC1F1(1)_5-79-36 | 2 | AAbbdd | 54 | 36.97531 | 15.75683 | 3.466636 | 6.133608 |
| YLD12BC1F1(1)_5-79-37 | 2 | AAbbdd | 45.66667 | 44.81752 | 17.58987 | 3.683895 | 6.520873 |
| YLD12BC1F1(1)_5-79-38 | 2 | AAbbdd | 51 | 38.75817 | 15.98971 | 3.414006 | 6.381545 |
| YLD12BC1F1(1)_5-79-39 | 2 | AAbbdd | 40 | 38.33333 | 15.53469 | 3.422312 | 6.291494 |
| YLD12BC1F1(1)_5-79-6 | 2 | AAbbdd | 51 | 36.66667 | 15.36857 | 3.386859 | 6.116934 |
| YLD12BC1F1(1)_5-79-9 | 2 | AAbbdd | 46.33333 | 39.85612 | 15.96989 | 3.486623 | 6.225975 |
| YLD12BC1F1(1)_5-96-12 | 2 | AAbbdd | 47.66667 | 43.00699 | 17.05406 | 3.617988 | 6.396201 |
| YLD12BC1F1(1)_5-96-13 | 2 | AAbbdd | 47.66667 | 35.1049 | 15.32359 | 3.340606 | 6.269469 |
| YLD12BC1F1(1)_5-96-14 | 2 | AAbbdd | 50 | 40.53333 | 16.68316 | 3.56416 | 6.347527 |
| YLD12BC1F1(1)_5-96-18 | 2 | AAbbdd | 56.66667 | 46.23529 | 17.58373 | 3.704524 | 6.391765 |
| YLD12BC1F1(1)_5-96-19 | 2 | AAbbdd | 65 | 47.4359 | 18.3186 | 3.81716 | 6.499781 |
| YLD12BC1F1(1)_5-96-20 | 2 | AAbbdd | 57 | 43.85965 | 17.2159 | 3.633849 | 6.386976 |
| YLD12BC1F1(1)_5-96-23 | 2 | AAbbdd | 55.66667 | 46.52695 | 17.94326 | 3.756868 | 6.420112 |
| YLD12BC1F1(1)_5-96-24 | 2 | AAbbdd | 43.33333 | 42.84615 | 17.00483 | 3.608704 | 6.408619 |
| YLD12BC1F1(1)_5-96-26 | 2 | AAbbdd | 47.66667 | 50.62937 | 19.03145 | 3.840358 | 6.710438 |
| YLD12BC1F1(1)_5-96-30 | 2 | AAbbdd | 30.66667 | 53.15217 | 19.71097 | 3.838208 | 7.010562 |
| YLD12BC1F1(1)_5-96-31 | 2 | AAbbdd | 54 | 41.35802 | 16.38569 | 3.515999 | 6.269969 |
| YLD12BC1F1(1)_5-96-33 | 2 | AAbbdd | 58 | 42.12644 | 17.13461 | 3.652585 | 6.349062 |
| YLD12BC1F1(1)_5-96-35 | 2 | AAbbdd | 53.33333 | 40.5625 | 16.73227 | 3.573152 | 6.303904 |
| YLD12BC1F1(1)_5-96-37 | 2 | AAbbdd | 55.33333 | 47.6506 | 18.30322 | 3.755762 | 6.59354 |
| YLD12BC1F1(1)_5-96-38 | 2 | AAbbdd | 48 | 38.125 | 15.85017 | 3.450877 | 6.328889 |
| YLD12BC1F1(1)_5-96-5 | 2 | AAbbdd | 56 | 43.63095 | 17.33488 | 3.639959 | 6.464763 |
| YLD12BC1F1(1)_5-96-6 | 2 | AAbbdd | 50 | 40.2 | 16.2748 | 3.517831 | 6.267883 |
| YLD12BC1F1(1)_5-144-1 | 4 | aaBBDD | 47.66667 | 38.32168 | 15.84088 | 3.446142 | 6.174463 |
| YLD12BC1F1(1)_5-144-10 | 4 | aaBBDD | 50 | 35.93333 | 15.28143 | 3.34336 | 6.318768 |
| YLD12BC1F1(1)_5-144-2 | 4 | aaBBDD | 44.66667 | 39.02985 | 16.07593 | 3.444518 | 6.30497 |
| YLD12BC1F1(1)_5-144-3 | 4 | aaBBDD | 43.33333 | 41.30769 | 16.71952 | 3.51915 | 6.434432 |
| YLD12BC1F1(1)_5-144-4 | 4 | aaBBDD | 47.66667 | 40.76923 | 16.53205 | 3.525961 | 6.406898 |
| YLD12BC1F1(1)_5-144-5 | 4 | aaBBDD | 53 | 39.74843 | 16.46057 | 3.538855 | 6.268282 |
| YLD12BC1F1(1)_5-144-6 | 4 | aaBBDD | 43 | 44.4186 | 17.46591 | 3.654382 | 6.580093 |
| YLD12BC1F1(1)_5-144-7 | 4 | aaBBDD | 54.66667 | 41.95122 | 16.94782 | 3.587006 | 6.417595 |
| YLD12BC1F1(1)_5-144-8 | 4 | aaBBDD | 49 | 34.35374 | 14.83691 | 3.267048 | 6.159366 |
| YLD12BC1F1(1)_5-144-9 | 4 | aaBBDD | 43.66667 | 39.69466 | 16.33669 | 3.509306 | 6.374115 |
| YLD12BC1F1(1)_5-84-1 | 4 | aaBBDD | 46.66667 | 48.21429 | 18.07014 | 3.851795 | 6.326996 |
| YLD12BC1F1(1)_5-84-10 | 4 | aaBBDD | 57.33333 | 45.5814 | 17.67827 | 3.71826 | 6.394955 |
| YLD12BC1F1(1)_5-84-2 | 4 | aaBBDD | 52.66667 | 40.82278 | 16.41905 | 3.566873 | 6.20122 |
| YLD12BC1F1(1)_5-84-3 | 4 | aaBBDD | 36 | 48.7037 | 18.16002 | 3.749625 | 6.547445 |
| YLD12BC1F1(1)_5-84-4 | 4 | aaBBDD | 56.33333 | 47.27811 | 18.26235 | 3.761675 | 6.529586 |
| YLD12BC1F1(1)_5-84-5 | 4 | aaBBDD | 63.33333 | 42.78947 | 17.00456 | 3.612895 | 6.30867 |
| YLD12BC1F1(1)_5-84-6 | 4 | aaBBDD | 46 | 52.46377 | 19.3325 | 3.942982 | 6.688573 |
| YLD12BC1F1(1)_5-84-7 | 4 | aaBBDD | 48.33333 | 49.37931 | 18.63975 | 3.787495 | 6.624022 |
| YLD12BC1F1(1)_5-84-8 | 4 | aaBBDD | 53.33333 | 36.3125 | 15.36595 | 3.384423 | 6.163012 |
| YLD12BC1F1(1)_5-84-9 | 4 | aaBBDD | 41.33333 | 46.20968 | 17.94755 | 3.724344 | 6.479948 |
| YLD12BC1F1(1)_1-9-1 | 4 | AAbbDD | 50.66667 | 46.11842 | 17.95823 | 3.687987 | 6.47581 |
| YLD12BC1F1(1)_1-9-10 | 4 | AAbbDD | 59.66667 | 46.75978 | 18.04557 | 3.74555 | 6.498929 |
| YLD12BC1F1(1)_1-9-2 | 4 | AAbbDD | 41.33333 | 53.22581 | 19.5662 | 3.90878 | 6.702263 |
| YLD12BC1F1(1)_1-9-4 | 4 | AAbbDD | 38.66667 | 60.86207 | 20.65103 | 4.104469 | 6.790242 |
| YLD12BC1F1(1)_1-9-5 | 4 | AAbbDD | 55.33333 | 46.92771 | 18.14109 | 3.752605 | 6.465271 |
| YLD12BC1F1(1)_1-9-7 | 4 | AAbbDD | 45 | 55.7037 | 19.70378 | 4.00481 | 6.674987 |
| YLD12BC1F1(1)_1-9-8 | 4 | AAbbDD | 53.33333 | 44.9375 | 17.1624 | 3.708756 | 6.147996 |
| YLD12BC1F1(1)_5-99-1 | 4 | AAbbDD | 52.33333 | 46.6242 | 17.93907 | 3.726569 | 6.474449 |
| YLD12BC1F1(1)_5-99-10 | 4 | AAbbDD | 56.66667 | 46.29412 | 18.04663 | 3.703669 | 6.574864 |
| YLD12BC1F1(1)_5-99-2 | 4 | AAbbDD | 47.66667 | 46.36364 | 17.923 | 3.756915 | 6.400307 |
| YLD12BC1F1(1)_5-99-3 | 4 | AAbbDD | 47 | 46.66667 | 17.90577 | 3.72121 | 6.486871 |
| YLD12BC1F1(1)_5-99-4 | 4 | AAbbDD | 52 | 45.83333 | 17.66844 | 3.724234 | 6.368928 |
| YLD12BC1F1(1)_5-99-5 | 4 | AAbbDD | 55 | 46.48485 | 17.82194 | 3.743151 | 6.344645 |
| YLD12BC1F1(1)_5-99-6 | 4 | AAbbDD | 39.33333 | 54.40678 | 19.53776 | 3.914196 | 6.776862 |
| YLD12BC1F1(1)_5-99-7 | 4 | AAbbDD | 52.33333 | 48.66242 | 18.41785 | 3.782532 | 6.514685 |
| YLD12BC1F1(1)_5-99-8 | 4 | AAbbDD | 55.33333 | 49.93976 | 18.31345 | 3.828425 | 6.409964 |
| YLD12BC1F1(1)_5-99-9 | 4 | AAbbDD | 52.66667 | 49.43038 | 19.0483 | 3.824657 | 6.875359 |
| YLD12BC1F1(1)_5-112-1 | 4 | AABBdd | 46 | 41.30435 | 16.41444 | 3.585984 | 6.196088 |
| YLD12BC1F1(1)_5-112-10 | 4 | AABBdd | 60.66667 | 45.76923 | 17.93506 | 3.754718 | 6.42781 |
| YLD12BC1F1(1)_5-112-2 | 4 | AABBdd | 42 | 47.22222 | 17.87156 | 3.740033 | 6.373183 |
| YLD12BC1F1(1)_5-112-3 | 4 | AABBdd | 46.66667 | 49.92857 | 18.5537 | 3.82152 | 6.549132 |
| YLD12BC1F1(1)_5-112-4 | 4 | AABBdd | 49.33333 | 44.52703 | 17.50706 | 3.675107 | 6.458022 |
| YLD12BC1F1(1)_5-112-5 | 4 | AABBdd | 47.66667 | 43.63636 | 17.15112 | 3.707776 | 6.212647 |
| YLD12BC1F1(1)_5-112-6 | 4 | AABBdd | 43 | 47.82946 | 17.86032 | 3.697344 | 6.564478 |
| YLD12BC1F1(1)_5-112-7 | 4 | AABBdd | 57.66667 | 45.72254 | 17.49991 | 3.747833 | 6.273877 |
| YLD12BC1F1(1)_5-112-8 | 4 | AABBdd | 38 | 42.19298 | 16.4321 | 3.605549 | 6.25713 |
| YLD12BC1F1(1)_5-112-9 | 4 | AABBdd | 44.33333 | 36.16541 | 15.62473 | 3.428214 | 6.237613 |
| YLD12BC1F1(1)_5-113-1 | 4 | AABBdd | 49.33333 | 42.2973 | 16.39543 | 3.560874 | 6.192522 |
| YLD12BC1F1(1)_5-113-10 | 4 | AABBdd | 56.33333 | 41.12426 | 16.61933 | 3.578538 | 6.291797 |
| YLD12BC1F1(1)_5-113-2 | 4 | AABBdd | 47.33333 | 46.05634 | 17.9078 | 3.721624 | 6.449829 |
| YLD12BC1F1(1)_5-113-3 | 4 | AABBdd | 43.66667 | 39.61832 | 16.16939 | 3.516066 | 6.278927 |
| YLD12BC1F1(1)_5-113-4 | 4 | AABBdd | 50 | 42.2 | 16.66524 | 3.593565 | 6.305424 |
| YLD12BC1F1(1)_5-113-5 | 4 | AABBdd | 52.66667 | 42.1519 | 16.82271 | 3.620075 | 6.244946 |
| YLD12BC1F1(1)_5-113-6 | 4 | AABBdd | 47 | 42.69504 | 17.12058 | 3.597245 | 6.564344 |
| YLD12BC1F1(1)_5-113-7 | 4 | AABBdd | 54 | 47.22222 | 18.19193 | 3.82645 | 6.424615 |
| YLD12BC1F1(1)_5-113-8 | 4 | AABBdd | 52 | 46.28205 | 17.62644 | 3.720255 | 6.385413 |
| YLD12BC1F1(1)_5-138-1 | 4 | AABBdd | 51.66667 | 46.83871 | 17.69097 | 3.763893 | 6.326814 |
| YLD12BC1F1(1)_5-138-10 | 4 | AABBdd | 53.66667 | 45.15528 | 18.09426 | 3.692508 | 6.709266 |
| YLD12BC1F1(1)_5-138-2 | 4 | AABBdd | 49.66667 | 45.57047 | 17.51632 | 3.726988 | 6.313653 |
| YLD12BC1F1(1)_5-138-3 | 4 | AABBdd | 49.66667 | 46.6443 | 17.76072 | 3.728238 | 6.427521 |
| YLD12BC1F1(1)_5-138-4 | 4 | AABBdd | 54.33333 | 45.03067 | 17.68259 | 3.658332 | 6.564678 |
| YLD12BC1F1(1)_5-138-5 | 4 | AABBdd | 45.66667 | 47.29927 | 18.09845 | 3.817083 | 6.419425 |
| YLD12BC1F1(1)_5-138-6 | 4 | AABBdd | 47.33333 | 48.23944 | 18.01516 | 3.778734 | 6.486812 |
| YLD12BC1F1(1)_5-138-7 | 4 | AABBdd | 55.66667 | 47.72455 | 18.17074 | 3.794233 | 6.458875 |
| YLD12BC1F1(1)_5-138-8 | 4 | AABBdd | 46 | 47.6087 | 18.00333 | 3.760034 | 6.416157 |
| YLD12BC1F1(1)_5-138-9 | 4 | AABBdd | 50.66667 | 46.71053 | 17.76118 | 3.767629 | 6.406134 |
| YLD12BC1F1(1)_5-147-1 | 4 | AABBdd | 48.33333 | 44.13793 | 17.32273 | 3.607109 | 6.489015 |
| YLD12BC1F1(1)_5-147-10 | 4 | AABBdd | 59.66667 | 41.67598 | 17.01853 | 3.598905 | 6.387867 |
| YLD12BC1F1(1)_5-147-2 | 4 | AABBdd | 54 | 47.83951 | 17.93418 | 3.795599 | 6.297174 |
| YLD12BC1F1(1)_5-147-3 | 4 | AABBdd | 40 | 53.25 | 20.26103 | 3.910185 | 7.056073 |
| YLD12BC1F1(1)_5-147-4 | 4 | AABBdd | 48.66667 | 49.24658 | 18.91247 | 3.788685 | 6.805305 |
| YLD12BC1F1(1)_5-147-5 | 4 | AABBdd | 51.33333 | 46.55844 | 18.30511 | 3.771988 | 6.686796 |
| YLD12BC1F1(1)_5-147-6 | 4 | AABBdd | 55.66667 | 40.83832 | 16.7471 | 3.560624 | 6.449253 |
| YLD12BC1F1(1)_5-147-7 | 4 | AABBdd | 41.33333 | 40.96774 | 16.78735 | 3.508352 | 6.656472 |
| YLD12BC1F1(1)_5-147-8 | 4 | AABBdd | 43 | 42.55814 | 17.04139 | 3.618631 | 6.378317 |
| YLD12BC1F1(1)_5-147-9 | 4 | AABBdd | 39.33333 | 48.72881 | 18.80071 | 3.781933 | 6.785303 |
| YLD12BC1F1(1)_1-54-1 | 6 | AABBDD | 45 | 43.85185 | 17.02931 | 3.667278 | 6.275397 |
| YLD12BC1F1(1)_1-54-10 | 6 | AABBDD | 46 | 49.56522 | 18.60001 | 3.837337 | 6.559144 |
| YLD12BC1F1(1)_1-54-2 | 6 | AABBDD | 52.33333 | 41.01911 | 16.7416 | 3.608211 | 6.282545 |
| YLD12BC1F1(1)_1-54-3 | 6 | AABBDD | 43.33333 | 45.30769 | 17.83696 | 3.685594 | 6.655219 |
| YLD12BC1F1(1)_1-54-4 | 6 | AABBDD | 53 | 43.14465 | 17.01972 | 3.63552 | 6.260283 |
| YLD12BC1F1(1)_1-54-5 | 6 | AABBDD | 49 | 42.2449 | 17.13719 | 3.621462 | 6.470648 |
| YLD12BC1F1(1)_1-54-6 | 6 | AABBDD | 44 | 46.28788 | 17.77815 | 3.774866 | 6.406806 |
| YLD12BC1F1(1)_1-54-7 | 6 | AABBDD | 51 | 44.57516 | 17.37582 | 3.739091 | 6.214347 |
| YLD12BC1F1(1)_1-54-8 | 6 | AABBDD | 41.33333 | 42.33871 | 17.30307 | 3.602254 | 6.506837 |
| YLD12BC1F1(1)_1-54-9 | 6 | AABBDD | 53 | 42.38994 | 16.96889 | 3.673798 | 6.273819 |
| YLD12BC1F1(1)_5-40-1 | 6 | AABBDD | 56.33333 | 42.95858 | 16.8997 | 3.670786 | 6.1389 |
| YLD12BC1F1(1)_5-40-10 | 6 | AABBDD | 51.33333 | 44.74026 | 17.23369 | 3.688303 | 6.280989 |
| YLD12BC1F1(1)_5-40-2 | 6 | AABBDD | 51.33333 | 44.09091 | 17.39672 | 3.719106 | 6.32449 |
| YLD12BC1F1(1)_5-40-3 | 6 | AABBDD | 31.66667 | 46.21053 | 18.26841 | 3.713288 | 6.724859 |
| YLD12BC1F1(1)_5-40-4 | 6 | AABBDD | 52.66667 | 43.29114 | 16.9505 | 3.629783 | 6.288028 |
| YLD12BC1F1(1)_5-40-5 | 6 | AABBDD | 50 | 41 | 16.59201 | 3.608532 | 6.211522 |
| YLD12BC1F1(1)_5-40-6 | 6 | AABBDD | 50 | 47.2 | 18.38609 | 3.811507 | 6.593201 |
| YLD12BC1F1(1)_5-40-7 | 6 | AABBDD | 51 | 46.79739 | 17.83087 | 3.806115 | 6.277752 |
| YLD12BC1F1(1)_5-40-8 | 6 | AABBDD | 40.66667 | 42.54098 | 16.7007 | 3.656484 | 6.202542 |
| YLD12BC1F1(1)_5-40-9 | 6 | AABBDD | 47.66667 | 44.8951 | 17.70437 | 3.688434 | 6.527089 |
| YLD12BC1F1(1)_5-9-1 | 6 | AABBDD | 36.33333 | 46.6055 | 18.16751 | 3.811756 | 6.432502 |
| YLD12BC1F1(1)_5-9-10 | 6 | AABBDD | 34.66667 | 46.25 | 18.12465 | 3.78372 | 6.518269 |
| YLD12BC1F1(1)_5-9-2 | 6 | AABBDD | 34.66667 | 44.32692 | 17.06011 | 3.697485 | 6.218754 |
| YLD12BC1F1(1)_5-9-4 | 6 | AABBDD | 50.33333 | 44.50331 | 17.59096 | 3.73057 | 6.396677 |
| YLD12BC1F1(1)_5-9-5 | 6 | AABBDD | 56.66667 | 45.94118 | 18.12981 | 3.75862 | 6.613621 |
| YLD12BC1F1(1)_5-9-6 | 6 | AABBDD | 33.66667 | 49.10891 | 18.72735 | 3.819316 | 6.681665 |
| YLD12BC1F1(1)_5-9-7 | 6 | AABBDD | 39 | 47.17949 | 18.14215 | 3.797579 | 6.486753 |
| YLD12BC1F1(1)_5-9-8 | 6 | AABBDD | 30.33333 | 46.37363 | 18.12132 | 3.765444 | 6.509713 |
| YLD12BC1F1(1)_5-9-9 | 6 | AABBDD | 39.33333 | 45.25424 | 17.92807 | 3.714856 | 6.579633 |

**Supplementary Table 4.** Phenotypic data collected for T5 population derived from C538-1.

| ID | Dosage | genotype | SeedNum | TGW | GA | GW | GL |
| --- | --- | --- | --- | --- | --- | --- | --- |
| C538-1-78-2-22-11-11 | 0 | aabbdd | 32.3333333 | 37.5257732 | 14.5922076 | 3.42828541 | 5.73516563 |
| C538-1-78-2-22-11-20 | 0 | aabbdd | 40 | 40.1666667 | 15.1312173 | 3.54944194 | 5.66241668 |
| C538-1-78-2-22-11-22 | 0 | aabbdd | 36 | 40.6481481 | 15.2664728 | 3.5885002 | 5.67103843 |
| C538-1-78-2-22-11-26 | 0 | aabbdd | 51.6666667 | 41.3548387 | 15.7010366 | 3.62072443 | 5.7451058 |
| C538-1-78-2-22-16-1 | 0 | aabbdd | 44 | 40.1515152 | 15.245967 | 3.55343471 | 5.69293701 |
| C538-1-78-2-22-16-10 | 0 | aabbdd | 33 | 39.7979798 | 15.4910839 | 3.51320484 | 5.82929732 |
| C538-1-78-2-22-16-2 | 0 | aabbdd | 38.3333333 | 40.3478261 | 15.4384359 | 3.60837133 | 5.72139585 |
| C538-1-78-2-22-16-3 | 0 | aabbdd | 50.6666667 | 39.2763158 | 15.2164382 | 3.53449345 | 5.75090294 |
| C538-1-78-2-22-16-4 | 0 | aabbdd | 42.3333333 | 38.5826772 | 15.1236614 | 3.50762207 | 5.76372953 |
| C538-1-78-2-22-16-5 | 0 | aabbdd | 58.3333333 | 40.4571429 | 15.520903 | 3.60949775 | 5.70091421 |
| C538-1-78-2-22-16-6 | 0 | aabbdd | 52.6666667 | 39.4303797 | 15.2391173 | 3.57176121 | 5.66168599 |
| C538-1-78-2-22-16-7 | 0 | aabbdd | 47 | 41.2056738 | 15.4649526 | 3.63769586 | 5.67847917 |
| C538-1-78-2-22-16-8 | 0 | aabbdd | 39 | 37.3504274 | 14.8062777 | 3.45965797 | 5.67289517 |
| C538-1-78-2-22-16-9 | 0 | aabbdd | 41 | 38.3739837 | 15.0778845 | 3.50658029 | 5.6870059 |
| C538-1-78-2-22-27-1 | 0 | aabbdd | 51.3333333 | 39.4155844 | 15.0802878 | 3.56780019 | 5.64031424 |
| C538-1-78-2-22-27-10 | 0 | aabbdd | 42.6666667 | 39.765625 | 15.2074156 | 3.5087696 | 5.75246414 |
| C538-1-78-2-22-27-2 | 0 | aabbdd | 49 | 40.8843537 | 15.5864452 | 3.645339 | 5.72908776 |
| C538-1-78-2-22-27-3 | 0 | aabbdd | 26.6666667 | 41.75 | 15.8126052 | 3.60513154 | 5.87691356 |
| C538-1-78-2-22-27-4 | 0 | aabbdd | 35 | 37.3333333 | 14.6828446 | 3.47724093 | 5.6615572 |
| C538-1-78-2-22-27-5 | 0 | aabbdd | 50.6666667 | 41.1184211 | 15.3872346 | 3.62242791 | 5.63890824 |
| C538-1-78-2-22-27-7 | 0 | aabbdd | 43.6666667 | 40.5343511 | 15.2212647 | 3.61994445 | 5.60603159 |
| C538-1-78-2-22-27-8 | 0 | aabbdd | 24 | 40.9722222 | 15.4748079 | 3.50640444 | 5.88107133 |
| C538-1-78-2-22-27-9 | 0 | aabbdd | 42 | 40.4761905 | 15.2154384 | 3.58774544 | 5.6044953 |
| C538-1-78-2-22-37-1 | 0 | aabbdd | 45 | 38.2222222 | 14.6986162 | 3.50368981 | 5.61940656 |
| C538-1-78-2-22-37-2 | 0 | aabbdd | 37.6666667 | 37.699115 | 14.6557994 | 3.48438039 | 5.58257101 |
| C538-1-78-2-22-37-4 | 0 | aabbdd | 37 | 41.3513514 | 15.769558 | 3.6061526 | 5.78035159 |
| C538-1-78-2-22-37-5 | 0 | aabbdd | 49.6666667 | 38.9261745 | 15.0726814 | 3.53264997 | 5.64862277 |
| C538-1-78-2-22-37-6 | 0 | aabbdd | 43.6666667 | 34.5038168 | 14.254882 | 3.39729631 | 5.58179233 |
| C538-1-78-2-22-37-8 | 0 | aabbdd | 28.3333333 | 38.4705882 | 14.9202071 | 3.54576337 | 5.57609965 |
| C538-1-78-2-22-37-9 | 0 | aabbdd | 47.3333333 | 41.2676056 | 15.7394819 | 3.58905861 | 5.84627296 |
| C538-1-78-2-22-41-13 | 0 | aabbdd | 27 | 37.7777778 | 14.3242624 | 3.49461714 | 5.48734034 |
| C538-1-78-2-22-41-15 | 0 | aabbdd | 24.3333333 | 38.2191781 | 14.8916775 | 3.48367074 | 5.70182055 |
| C538-1-78-2-22-46-13 | 0 | aabbdd | 41 | 38.699187 | 15.0333766 | 3.50620189 | 5.7301531 |
| C538-1-78-2-22-46-16 | 0 | aabbdd | 55.6666667 | 39.1616766 | 15.0054885 | 3.51028326 | 5.94692068 |
| C538-1-78-2-22-46-22 | 0 | aabbdd | 44.3333333 | 40.1503759 | 15.2848294 | 3.53324503 | 5.73468935 |
| C538-1-78-2-22-46-27 | 0 | aabbdd | 47.6666667 | 39.020979 | 14.9748949 | 3.53832406 | 5.63254586 |
| C538-1-78-2-22-46-4 | 0 | aabbdd | 40.6666667 | 39.8360656 | 15.0404035 | 3.54360409 | 5.70473175 |
| C538-1-78-2-22-47-10 | 0 | aabbdd | 42.6666667 | 40.78125 | 15.2291865 | 3.52469235 | 5.73548805 |
| C538-1-78-2-22-47-11 | 0 | aabbdd | 47.3333333 | 40.6338028 | 15.356994 | 3.57908733 | 5.85574959 |
| C538-1-78-2-22-47-14 | 0 | aabbdd | 55 | 38.9090909 | 14.990426 | 3.51952 | 5.67534554 |
| C538-1-78-2-22-47-17 | 0 | aabbdd | 44.3333333 | 38.1954887 | 15.1510715 | 3.54049301 | 5.77115941 |
| C538-1-78-2-22-47-20 | 0 | aabbdd | 31 | 38.6021505 | 15.0901131 | 3.52152451 | 5.68898223 |
| C538-1-78-2-22-47-24 | 0 | aabbdd | 39 | 39.7435897 | 14.9534868 | 3.51406886 | 5.60204421 |
| C538-1-78-2-22-47-25 | 0 | aabbdd | 44.6666667 | 41.4179104 | 15.491359 | 3.60313038 | 5.69247204 |
| C538-1-78-2-22-47-29 | 0 | aabbdd | 44.3333333 | 40.9022556 | 15.62845 | 3.57704595 | 5.79163905 |
| C538-1-78-2-22-47-6 | 0 | aabbdd | 35 | 38.1904762 | 14.4178746 | 3.48866346 | 5.51539287 |
| C538-1-78-2-22-76-2 | 0 | aabbdd | 47.3333333 | 40.7746479 | 15.4729052 | 3.60897821 | 5.71623739 |
| C538-1-78-2-22-76-3 | 0 | aabbdd | 50.3333333 | 37.0860927 | 14.9692465 | 3.44474439 | 5.74942084 |
| C538-1-78-2-22-76-7 | 0 | aabbdd | 32 | 43.75 | 16.2242584 | 3.77379871 | 5.68297174 |
| C538-1-78-2-22-76-8 | 0 | aabbdd | 29.3333333 | 43.9772727 | 16.2960451 | 3.70469599 | 5.83475508 |
| C538-1-78-2-22-76-9 | 0 | aabbdd | 30.6666667 | 41.5217391 | 16.197425 | 3.7121264 | 5.80976314 |
| C538-1-78-2-22-41-10 | 1 | aabbDd | 45.6666667 | 41.3138686 | 15.707762 | 3.56829055 | 5.85494502 |
| C538-1-78-2-22-41-12 | 1 | aabbDd | 51 | 39.8039216 | 15.4182724 | 3.54469405 | 5.82279446 |
| C538-1-78-2-22-41-3 | 1 | aabbDd | 45.3333333 | 41.3235294 | 15.7140008 | 3.5854513 | 5.80722137 |
| C538-1-78-2-22-41-4 | 1 | aabbDd | 46 | 41.4492754 | 15.6928197 | 3.54379801 | 5.8772233 |
| C538-1-78-2-22-41-6 | 1 | aabbDd | 24 | 38.75 | 15.0456943 | 3.4846754 | 5.85037967 |
| C538-1-78-2-22-41-8 | 1 | aabbDd | 35 | 38.5714286 | 15.0602234 | 3.43259159 | 5.85060515 |
| C538-1-78-2-22-11-1 | 1 | aaBbdd | 52.3333333 | 40.5095541 | 15.5268462 | 3.55332222 | 5.8416216 |
| C538-1-78-2-22-11-12 | 1 | aaBbdd | 57 | 34.9122807 | 14.5207475 | 3.29966903 | 5.96804781 |
| C538-1-78-2-22-11-14 | 1 | aaBbdd | 50.3333333 | 40.5960265 | 15.538191 | 3.56390693 | 5.78133892 |
| C538-1-78-2-22-11-17 | 1 | aaBbdd | 52.6666667 | 37.278481 | 14.744422 | 3.43678321 | 5.77470421 |
| C538-1-78-2-22-11-2 | 1 | aaBbdd | 43.6666667 | 39.9236641 | 15.4632892 | 3.47378499 | 5.9361555 |
| C538-1-78-2-22-11-23 | 1 | aaBbdd | 35.3333333 | 40.3773585 | 15.3789739 | 3.58810391 | 5.72668922 |
| C538-1-78-2-22-11-24 | 1 | aaBbdd | 41 | 39.8373984 | 15.4012065 | 3.51672769 | 5.82471415 |
| C538-1-78-2-22-11-25 | 1 | aaBbdd | 44 | 40.530303 | 15.5471761 | 3.56052265 | 5.77182354 |
| C538-1-78-2-22-11-27 | 1 | aaBbdd | 45.6666667 | 42.919708 | 16.0912658 | 3.63126422 | 5.86011053 |
| C538-1-78-2-22-11-28 | 1 | aaBbdd | 41 | 37.5609756 | 15.0549364 | 3.46972402 | 5.79670816 |
| C538-1-78-2-22-11-3 | 1 | aaBbdd | 37 | 36.8468468 | 14.8325665 | 3.4060982 | 5.81776312 |
| C538-1-78-2-22-11-30 | 1 | aaBbdd | 32.3333333 | 39.3814433 | 15.5272788 | 3.55704625 | 5.87501974 |
| C538-1-78-2-22-11-4 | 1 | aaBbdd | 49.6666667 | 39.8657718 | 15.5194308 | 3.55223334 | 5.7990871 |
| C538-1-78-2-22-11-5 | 1 | aaBbdd | 45 | 41.1851852 | 15.7455019 | 3.5969479 | 5.92072453 |
| C538-1-78-2-22-11-6 | 1 | aaBbdd | 35.3333333 | 41.3207547 | 15.5715652 | 3.56954811 | 5.77107241 |
| C538-1-78-2-22-11-7 | 1 | aaBbdd | 31.6666667 | 41.0526316 | 15.7410932 | 3.52996193 | 5.95994157 |
| C538-1-78-2-22-11-8 | 1 | aaBbdd | 36.3333333 | 38.2568807 | 15.0987033 | 3.45249882 | 5.85909171 |
| C538-1-78-2-22-30-20 | 1 | aaBbdd | 40.3333333 | 39.6694215 | 15.2678954 | 3.49839602 | 5.81277477 |
| C538-1-78-2-22-30-24 | 1 | aaBbdd | 35.6666667 | 40.1869159 | 15.2906932 | 3.55240518 | 5.76882191 |
| C538-1-78-2-22-30-34 | 1 | aaBbdd | 39 | 39.4017094 | 15.3857409 | 3.52578041 | 5.81220011 |
| C538-1-78-2-22-30-36 | 1 | aaBbdd | 35 | 42.6666667 | 16.1456156 | 3.62302009 | 5.9598797 |
| C538-1-78-2-22-30-55 | 1 | aaBbdd | 44.3333333 | 43.0075188 | 15.9528753 | 3.62670253 | 5.89009458 |
| C538-1-78-2-22-30-9 | 1 | aaBbdd | 49.6666667 | 41.2751678 | 15.6843149 | 3.55695478 | 5.87609038 |
| C538-1-78-2-22-46-1 | 1 | Aabbdd | 51 | 41.7647059 | 16.0058662 | 3.5772355 | 5.96645029 |
| C538-1-78-2-22-46-11 | 1 | Aabbdd | 37 | 40.1801802 | 15.4344438 | 3.49032747 | 5.85869516 |
| C538-1-78-2-22-46-2 | 1 | Aabbdd | 52 | 40.7692308 | 15.6915496 | 3.53624602 | 5.90466513 |
| C538-1-78-2-22-46-20 | 1 | Aabbdd | 51 | 40.9803922 | 15.7182677 | 3.56376288 | 5.84683862 |
| C538-1-78-2-22-46-21 | 1 | Aabbdd | 41 | 40.9756098 | 15.5731723 | 3.56565768 | 5.79952621 |
| C538-1-78-2-22-46-24 | 1 | Aabbdd | 37.6666667 | 38.9380531 | 15.1769395 | 3.48684191 | 5.81143907 |
| C538-1-78-2-22-46-28 | 1 | Aabbdd | 41 | 40.8943089 | 16.1029556 | 3.5340695 | 6.0594549 |
| C538-1-78-2-22-46-29 | 1 | Aabbdd | 45.3333333 | 40.4411765 | 15.9476561 | 3.53356904 | 6.00045415 |
| C538-1-78-2-22-46-30 | 1 | Aabbdd | 56.3333333 | 41.2426036 | 16.2083228 | 3.56807075 | 6.03842659 |
| C538-1-78-2-22-46-8 | 1 | Aabbdd | 36.6666667 | 35.8181818 | 14.621157 | 3.35180673 | 5.84544017 |
| C538-1-78-2-22-47-1 | 1 | Aabbdd | 58.6666667 | 39.4886364 | 15.5577999 | 3.52508753 | 5.86485968 |
| C538-1-78-2-22-47-12 | 1 | Aabbdd | 42.6666667 | 41.484375 | 15.7258447 | 3.59826895 | 5.88568677 |
| C538-1-78-2-22-47-13 | 1 | Aabbdd | 46.3333333 | 38.5611511 | 14.9303003 | 3.45625002 | 5.74791178 |
| C538-1-78-2-22-47-16 | 1 | Aabbdd | 46.3333333 | 39.7841727 | 15.1494061 | 3.50237479 | 5.86764392 |
| C538-1-78-2-22-47-18 | 1 | Aabbdd | 46.6666667 | 38.9285714 | 15.1231912 | 3.46862125 | 5.82038229 |
| C538-1-78-2-22-47-19 | 1 | Aabbdd | 42.3333333 | 39.2913386 | 15.2379032 | 3.52592326 | 5.73881353 |
| C538-1-78-2-22-47-2 | 1 | Aabbdd | 52.3333333 | 40.5732484 | 15.5603986 | 3.56141638 | 5.82159454 |
| C538-1-78-2-22-47-22 | 1 | Aabbdd | 51.3333333 | 41.3636364 | 15.6239062 | 3.57223832 | 5.81698202 |
| C538-1-78-2-22-47-27 | 1 | Aabbdd | 40.3333333 | 41.1570248 | 15.7833307 | 3.57820501 | 5.86309431 |
| C538-1-78-2-22-47-3 | 1 | Aabbdd | 52.3333333 | 39.8089172 | 15.3231845 | 3.51404374 | 5.81089952 |
| C538-1-78-2-22-47-4 | 1 | Aabbdd | 33.6666667 | 41.1881188 | 15.471898 | 3.60818723 | 5.72616444 |
| C538-1-78-2-22-47-8 | 1 | Aabbdd | 43 | 39.1472868 | 15.5079834 | 3.4781924 | 5.93387224 |
| C538-1-78-2-22-47-9 | 1 | Aabbdd | 50 | 38.7333333 | 15.156657 | 3.48636022 | 5.80195407 |
| C538-1-78-2-22-41-1 | 2 | aabbDD | 30.6666667 | 39.3478261 | 15.1519317 | 3.46919838 | 5.83752204 |
| C538-1-78-2-22-41-11 | 2 | aabbDD | 39 | 40.5128205 | 15.6223052 | 3.53147473 | 5.88876696 |
| C538-1-78-2-22-41-7 | 2 | aabbDD | 32.6666667 | 39.2857143 | 15.2461977 | 3.51207494 | 5.82769122 |
| C538-1-78-2-22-14-29 | 2 | aaBbDd | 30.3333333 | 37.4725275 | 14.6718005 | 3.35083388 | 5.99540751 |
| C538-1-78-2-22-30-25 | 2 | aaBbDd | 37.6666667 | 40.2654867 | 15.319893 | 3.52112311 | 5.76615587 |
| C538-1-78-2-22-30-54 | 2 | aaBbDd | 60.6666667 | 41.7032967 | 15.9382528 | 3.60356414 | 5.88123395 |
| C538-1-78-2-22-11-16 | 2 | aaBBdd | 46.6666667 | 36.6428571 | 14.5457579 | 3.42182235 | 5.64007118 |
| C538-1-78-2-22-14-2 | 2 | aaBBdd | 39.3333333 | 36.1016949 | 14.2247849 | 3.29954014 | 5.77436204 |
| C538-1-78-2-22-14-23 | 2 | aaBBdd | 30 | 41.2222222 | 15.2452315 | 3.54091468 | 5.76949947 |
| C538-1-78-2-22-14-24 | 2 | aaBBdd | 50 | 38.2 | 15.0179823 | 3.47348328 | 5.72362227 |
| C538-1-78-2-22-14-6 | 2 | aaBBdd | 44.3333333 | 40.2255639 | 15.2696895 | 3.54052008 | 5.73570621 |
| C538-1-78-2-22-30-18 | 2 | AabbDd | 46.3333333 | 40.1438849 | 15.4629317 | 3.51900258 | 5.85348112 |
| C538-1-78-2-22-30-28 | 2 | AabbDd | 40.3333333 | 39.0082645 | 15.2441221 | 3.46863083 | 5.85434382 |
| C538-1-78-2-22-30-33 | 2 | AabbDd | 34 | 39.9019608 | 15.3613695 | 3.5441464 | 5.82169561 |
| C538-1-78-2-22-30-39 | 2 | AabbDd | 21.6666667 | 43.8461538 | 16.2177801 | 3.5967629 | 6.03567546 |
| C538-1-78-2-22-30-50 | 2 | AabbDd | 41.3333333 | 40.8870968 | 15.6284554 | 3.5627484 | 5.86367038 |
| C538-1-78-2-22-30-68 | 2 | AabbDd | 67 | 36.0199005 | 14.4429666 | 3.38037294 | 5.6265461 |
| C538-1-78-2-22-30-8 | 2 | AabbDd | 52.3333333 | 42.2929936 | 16.1440724 | 3.60224916 | 5.97554422 |
| C538-1-78-2-22-30-10 | 2 | AaBbdd | 52.6666667 | 43.0379747 | 16.0298332 | 3.613161 | 5.92213941 |
| C538-1-78-2-22-30-2 | 2 | AaBbdd | 41.6666667 | 39.84 | 15.8770225 | 3.56242899 | 6.00784532 |
| C538-1-78-2-22-30-40 | 2 | AaBbdd | 29 | 41.7241379 | 15.8741509 | 3.58246078 | 5.922902 |
| C538-1-78-2-22-30-48 | 2 | AaBbdd | 56 | 41.9047619 | 16.1788973 | 3.61159212 | 6.02628316 |
| C538-1-78-2-22-30-57 | 2 | AaBbdd | 36.6666667 | 43.3636364 | 16.2106264 | 3.61087877 | 5.97131696 |
| C538-1-78-2-22-30-61 | 2 | AaBbdd | 49.6666667 | 41.6107383 | 15.7932722 | 3.60749911 | 5.98940434 |
| C538-1-78-2-22-46-10 | 2 | AAbbdd | 36.6666667 | 42.0909091 | 16.2693836 | 3.59249391 | 6.03923559 |
| C538-1-78-2-22-46-12 | 2 | AAbbdd | 40.6666667 | 40.1639344 | 15.6227914 | 3.51032141 | 6.05551697 |
| C538-1-78-2-22-46-17 | 2 | AAbbdd | 40.3333333 | 42.9752066 | 15.897294 | 3.57361217 | 5.94273797 |
| C538-1-78-2-22-46-18 | 2 | AAbbdd | 39.6666667 | 40.6722689 | 15.5636285 | 3.53150499 | 5.94147513 |
| C538-1-78-2-22-46-19 | 2 | AAbbdd | 49.6666667 | 39.0604027 | 15.1339774 | 3.53042135 | 5.74613525 |
| C538-1-78-2-22-46-26 | 2 | AAbbdd | 45.3333333 | 42.0588235 | 16.0689454 | 3.57729005 | 5.93846445 |
| C538-1-78-2-22-46-3 | 2 | AAbbdd | 43 | 42.248062 | 16.1893575 | 3.62918178 | 5.97953517 |
| C538-1-78-2-22-46-5 | 2 | AAbbdd | 53.6666667 | 40.931677 | 15.5641966 | 3.53225668 | 5.83260631 |
| C538-1-78-2-22-46-6 | 2 | AAbbdd | 42 | 40.8730159 | 15.2242136 | 3.50886823 | 5.74055643 |
| C538-1-78-2-22-47-15 | 2 | AAbbdd | 44.6666667 | 39.0298507 | 15.2295293 | 3.52336907 | 5.76779896 |
| C538-1-78-2-22-30-19 | 3 | aaBbDD | 50 | 41.7333333 | 15.9375264 | 3.59363119 | 5.94979563 |
| C538-1-78-2-22-30-31 | 3 | aaBbDD | 40 | 42.4166667 | 15.9926158 | 3.58803709 | 5.93816999 |
| C538-1-78-2-22-30-49 | 3 | aaBbDD | 47.3333333 | 43.3098592 | 16.351703 | 3.67321343 | 6.00313151 |
| C538-1-78-2-22-14-10 | 3 | aaBBDd | 47.3333333 | 39.2957746 | 15.1425806 | 3.42733076 | 5.96996797 |
| C538-1-78-2-22-14-13 | 3 | aaBBDd | 40.3333333 | 43.3057851 | 16.3128814 | 3.63374211 | 6.00205217 |
| C538-1-78-2-22-14-17 | 3 | aaBBDd | 50 | 37.9333333 | 14.967882 | 3.47780838 | 5.78698881 |
| C538-1-78-2-22-14-18 | 3 | aaBBDd | 47.6666667 | 39.6503497 | 15.2269093 | 3.48848797 | 6.20719611 |
| C538-1-78-2-22-14-21 | 3 | aaBBDd | 50 | 40 | 15.4044018 | 3.55744411 | 5.77932768 |
| C538-1-78-2-22-14-28 | 3 | aaBBDd | 38.6666667 | 38.2758621 | 15.1933674 | 3.451945 | 5.90493444 |
| C538-1-78-2-22-30-35 | 3 | aaBBDd | 45 | 42 | 15.9685028 | 3.60826066 | 5.94051366 |
| C538-1-78-2-22-30-42 | 3 | aaBBDd | 43 | 41.7829457 | 16.0793955 | 3.62196233 | 5.94397436 |
| C538-1-78-2-22-30-51 | 3 | aaBBDd | 47 | 42.3404255 | 16.1191942 | 3.61540485 | 6.01704166 |
| C538-1-78-2-22-30-52 | 3 | AabbDD | 46.3333333 | 40.3597122 | 15.354315 | 3.49530406 | 5.89710221 |
| C538-1-78-2-22-30-29 | 3 | AaBbDd | 26.6666667 | 38.5 | 15.2734728 | 3.50669511 | 5.77977642 |
| C538-1-78-2-22-30-58 | 3 | AaBbDd | 42.6666667 | 40.703125 | 15.5384742 | 3.5075562 | 5.97110442 |
| C538-1-78-2-22-30-67 | 3 | AaBbDd | 57.3333333 | 41.627907 | 15.9627264 | 3.59897387 | 5.90614701 |
| C538-1-78-2-22-30-7 | 3 | AaBbDd | 48 | 41.3194444 | 15.8471142 | 3.54181112 | 5.98859469 |
| C538-1-78-2-22-30-69 | 3 | AaBBdd | 31.3333333 | 39.0425532 | 15.2722248 | 3.47145395 | 5.86448745 |
| C538-1-78-2-22-30-43 | 3 | AAbbDd | 36.6666667 | 40.6363636 | 15.6322883 | 3.52672503 | 5.94994453 |
| C538-1-78-2-22-30-5 | 3 | AAbbDd | 42.6666667 | 40.390625 | 15.5676894 | 3.54339788 | 5.87205161 |
| C538-1-78-2-22-14-16 | 4 | aaBBDD | 48.3333333 | 40.137931 | 15.3954249 | 3.53331515 | 5.75958677 |
| C538-1-78-2-22-14-20 | 4 | aaBBDD | 55.6666667 | 42.0958084 | 16.0790892 | 3.62291858 | 5.90540088 |
| C538-1-78-2-22-14-27 | 4 | aaBBDD | 38.3333333 | 40.5217391 | 15.7798726 | 3.53923906 | 5.89575612 |
| C538-1-78-2-22-14-9 | 4 | aaBBDD | 35.3333333 | 35.5660377 | 14.5273663 | 3.32998453 | 5.81506491 |
| C538-1-78-2-22-30-38 | 4 | aaBBDD | 30.3333333 | 43.4065934 | 16.5830613 | 3.66855661 | 6.04810367 |
| C538-1-78-2-22-62-10 | 4 | aaBBDD | 32 | 37.5 | 14.8374737 | 3.39054344 | 5.887501 |
| C538-1-78-2-22-62-11 | 4 | aaBBDD | 37.3333333 | 33.9285714 | 14.1867001 | 3.25525923 | 5.86075543 |
| C538-1-78-2-22-62-16 | 4 | aaBBDD | 45 | 37.3333333 | 14.8267302 | 3.42437919 | 5.8751348 |
| C538-1-78-2-22-62-24 | 4 | aaBBDD | 36 | 38.8888889 | 14.9094816 | 3.46873924 | 5.7562037 |
| C538-1-78-2-22-62-37 | 4 | aaBBDD | 37.3333333 | 40.3571429 | 15.9382467 | 3.51195226 | 6.11350826 |
| C538-1-78-2-22-62-5 | 4 | aaBBDD | 57.3333333 | 40.755814 | 15.5771801 | 3.55261849 | 5.81667014 |
| C538-1-78-2-22-62-9 | 4 | aaBBDD | 27.3333333 | 35.8536585 | 15.0356788 | 3.39070444 | 5.95807317 |
| C538-1-78-2-41-5-1 | 4 | aaBBDD | 37 | 41.2612613 | 15.4750981 | 3.51110018 | 5.89598593 |
| C538-1-78-2-41-5-10 | 4 | aaBBDD | 44 | 38.5606061 | 15.0637977 | 3.4706545 | 5.83893785 |
| C538-1-78-2-41-5-2 | 4 | aaBBDD | 42.6666667 | 38.90625 | 15.1131685 | 3.46789536 | 5.84478651 |
| C538-1-78-2-41-5-3 | 4 | aaBBDD | 53.6666667 | 32.2981366 | 13.9088837 | 3.26377523 | 5.7398898 |
| C538-1-78-2-41-5-4 | 4 | aaBBDD | 32.3333333 | 42.0618557 | 16.2975773 | 3.65512975 | 5.97068232 |
| C538-1-78-2-41-5-5 | 4 | aaBBDD | 54.3333333 | 39.202454 | 15.4022334 | 3.51259936 | 5.85338309 |
| C538-1-78-2-41-5-6 | 4 | aaBBDD | 30.3333333 | 39.4505495 | 15.5120518 | 3.60670493 | 5.77962759 |
| C538-1-78-2-41-5-7 | 4 | aaBBDD | 32.6666667 | 42.755102 | 15.9551589 | 3.56968813 | 5.91408851 |
| C538-1-78-2-41-5-8 | 4 | aaBBDD | 34 | 39.0196078 | 15.43064 | 3.50140604 | 5.83358083 |
| C538-1-78-2-41-5-9 | 4 | aaBBDD | 40 | 39.8333333 | 15.5147602 | 3.46038274 | 5.96393377 |
| C538-1-78-2-22-30-53 | 4 | AaBbDD | 53.6666667 | 42.9192547 | 16.1555354 | 3.62273283 | 5.92985999 |
| C538-1-78-2-22-30-59 | 4 | AaBbDD | 39 | 42.0512821 | 15.9551392 | 3.57923316 | 5.9346631 |
| C538-1-78-2-22-33-3 | 4 | AaBbDD | 33 | 37.7777778 | 15.0453424 | 3.33849785 | 5.98934543 |
| C538-1-78-2-22-62-12 | 4 | AaBbDD | 39.3333333 | 37.9661017 | 15.1280559 | 3.39790258 | 5.97502284 |
| C538-1-78-2-22-30-17 | 4 | AaBBDd | 41.6666667 | 42.4 | 15.8591878 | 3.57915572 | 5.90915789 |
| C538-1-78-2-22-30-4 | 4 | AaBBDd | 35.6666667 | 38.7850467 | 14.9771603 | 3.45565553 | 5.78843565 |
| C538-1-78-2-22-30-65 | 4 | AaBBDd | 39.6666667 | 41.3445378 | 15.6893275 | 3.57886454 | 5.85365635 |
| C538-1-78-2-22-33-29 | 4 | AAbbDD | 48.3333333 | 40.7586207 | 15.6293944 | 3.53860749 | 5.85908036 |
| C538-1-78-2-22-33-33 | 4 | AAbbDD | 38 | 40.9649123 | 15.6878829 | 3.46539534 | 5.98938838 |
| C538-1-78-2-22-33-34 | 4 | AAbbDD | 37.6666667 | 39.7345133 | 15.3921254 | 3.52051883 | 5.79551691 |
| C538-1-78-2-22-33-38 | 4 | AAbbDD | 45.3333333 | 40.8088235 | 15.8147997 | 3.55016208 | 6.02795563 |
| C538-1-78-2-22-33-6 | 4 | AAbbDD | 36 | 33.8888889 | 14.7559146 | 3.31952512 | 5.95190503 |
| C538-1-78-2-22-33-9 | 4 | AAbbDD | 38 | 34.0350877 | 14.6474912 | 3.29963588 | 5.92120601 |
| C538-1-78-2-22-30-16 | 4 | AABbDd | 38.6666667 | 39.8275862 | 15.4897814 | 3.47803041 | 5.93991203 |
| C538-1-78-2-22-30-47 | 4 | AABbDd | 40 | 41.1666667 | 15.8001701 | 3.56191597 | 5.97060912 |
| C538-1-78-2-22-30-6 | 4 | AABbDd | 47.6666667 | 40.979021 | 15.7332009 | 3.52832123 | 5.94882955 |
| C538-1-78-2-22-30-63 | 4 | AABbDd | 54.3333333 | 43.190184 | 16.5183388 | 3.66636584 | 6.09194738 |
| C538-1-78-2-22-30-11 | 4 | AABBdd | 36.3333333 | 42.6605505 | 15.6990888 | 3.57932001 | 5.85056245 |
| C538-1-78-2-22-30-64 | 4 | AABBdd | 43.6666667 | 42.1374046 | 16.0128022 | 3.57258898 | 6.14351318 |
| C538-1-78-2-22-62-13 | 5 | AaBBDD | 37.3333333 | 41.5178571 | 15.8091059 | 3.54919018 | 5.97408622 |
| C538-1-78-2-22-62-14 | 5 | AaBBDD | 42.3333333 | 38.3464567 | 14.9971743 | 3.43476728 | 5.84118837 |
| C538-1-78-2-22-62-15 | 5 | AaBBDD | 38.6666667 | 39.137931 | 15.1801934 | 3.44421707 | 6.40051251 |
| C538-1-78-2-22-62-22 | 5 | AaBBDD | 44.3333333 | 39.2481203 | 15.1701991 | 3.46590223 | 5.8619699 |
| C538-1-78-2-22-62-25 | 5 | AaBBDD | 40.3333333 | 41.9008264 | 15.9900843 | 3.57679899 | 5.97318501 |
| C538-1-78-2-22-62-28 | 5 | AaBBDD | 49 | 38.707483 | 15.1301319 | 3.47531682 | 5.87578838 |
| C538-1-78-2-22-62-32 | 5 | AaBBDD | 28.3333333 | 40.1176471 | 15.3549712 | 3.52697287 | 5.79655549 |
| C538-1-78-2-22-62-34 | 5 | AaBBDD | 40 | 39.25 | 15.2888295 | 3.46127538 | 5.89621762 |
| C538-1-78-2-22-62-35 | 5 | AaBBDD | 38.3333333 | 38.6086957 | 15.2648255 | 3.43845268 | 5.92044827 |
| C538-1-78-2-22-62-38 | 5 | AaBBDD | 42.3333333 | 38.503937 | 15.181844 | 3.44710692 | 5.92615331 |
| C538-1-78-2-22-62-40 | 5 | AaBBDD | 38.6666667 | 40.0862069 | 15.4521194 | 3.45491786 | 5.98723336 |
| C538-1-78-2-22-28-20 | 5 | AABbDD | 38 | 42.6315789 | 16.2107598 | 3.58955295 | 6.07631554 |
| C538-1-78-2-22-28-6 | 5 | AABbDD | 39 | 33.6752137 | 14.2517846 | 3.28321064 | 5.8729931 |
| C538-1-78-2-22-30-21 | 5 | AABbDD | 54.3333333 | 40.8588957 | 15.7250204 | 3.51899033 | 5.92135489 |
| C538-1-78-2-22-30-44 | 5 | AABbDD | 48.6666667 | 41.5068493 | 15.6679578 | 3.52373412 | 5.92951446 |
| C538-1-78-2-22-33-12 | 5 | AABbDD | 29 | 39.6551724 | 15.6672939 | 3.46041141 | 6.01932609 |
| C538-1-78-2-22-33-13 | 5 | AABbDD | 51.3333333 | 41.1038961 | 15.9195603 | 3.58759187 | 5.93083698 |
| C538-1-78-2-22-33-14 | 5 | AABbDD | 41 | 39.5934959 | 15.5121476 | 3.4909168 | 5.90615592 |
| C538-1-78-2-22-33-18 | 5 | AABbDD | 45 | 38.1481481 | 15.0826252 | 3.46858318 | 5.82063915 |
| C538-1-78-2-22-33-19 | 5 | AABbDD | 59 | 32.9378531 | 14.1446177 | 3.28009145 | 5.80004679 |
| C538-1-78-2-22-33-20 | 5 | AABbDD | 50.6666667 | 38.3552632 | 15.2972235 | 3.52424321 | 5.7747347 |
| C538-1-78-2-22-33-26 | 5 | AABbDD | 39.3333333 | 41.5254237 | 15.6030406 | 3.53356941 | 5.87939371 |
| C538-1-78-2-22-33-27 | 5 | AABbDD | 47 | 40.7092199 | 15.5431148 | 3.5777934 | 5.82956804 |
| C538-1-78-2-22-33-28 | 5 | AABbDD | 42 | 35.952381 | 14.8066398 | 3.40863994 | 5.80386375 |
| C538-1-78-2-22-33-32 | 5 | AABbDD | 38.3333333 | 40.7826087 | 15.4262418 | 3.54242968 | 5.82933829 |
| C538-1-78-2-22-33-35 | 5 | AABbDD | 38 | 41.0526316 | 15.7508867 | 3.50938738 | 5.94352166 |
| C538-1-78-2-22-33-4 | 5 | AABbDD | 49.3333333 | 42.2297297 | 16.258887 | 3.59690841 | 5.99547791 |
| C538-1-78-2-22-33-5 | 5 | AABbDD | 38.6666667 | 43.6206897 | 16.2791338 | 3.6025166 | 6.02658702 |
| C538-1-78-2-22-62-1 | 5 | AABbDD | 34 | 39.7058824 | 15.3606645 | 3.50146579 | 5.88882893 |
| C538-1-78-2-22-62-20 | 5 | AABbDD | 48 | 40.5555556 | 16.4239892 | 3.59583361 | 6.07201167 |
| C538-1-78-2-22-62-29 | 5 | AABbDD | 40 | 38.5833333 | 15.1183327 | 3.41882864 | 5.85000487 |
| C538-1-78-2-22-30-3 | 5 | AABBDd | 41 | 39.7560976 | 15.3594027 | 3.48380472 | 5.94124246 |
| C538-1-78-2-22-11-13 | 6 | AABBDD | 50.3333333 | 38.6092715 | 14.6973057 | 3.46545978 | 5.59647317 |
| C538-1-78-2-22-28-1 | 6 | AABBDD | 44 | 40.3030303 | 15.9355542 | 3.51760839 | 6.02615761 |
| C538-1-78-2-22-28-10 | 6 | AABBDD | 42.6666667 | 37.109375 | 14.9000242 | 3.41282131 | 5.84191106 |
| C538-1-78-2-22-28-11 | 6 | AABBDD | 53 | 40.3144654 | 15.8716887 | 3.51914497 | 6.0304628 |
| C538-1-78-2-22-28-12 | 6 | AABBDD | 42.3333333 | 38.8976378 | 15.3721762 | 3.47935216 | 5.89686315 |
| C538-1-78-2-22-28-13 | 6 | AABBDD | 47.6666667 | 37.6923077 | 14.9199534 | 3.45780606 | 5.73815973 |
| C538-1-78-2-22-28-15 | 6 | AABBDD | 44 | 41.3636364 | 15.5850401 | 3.56204592 | 5.78292631 |
| C538-1-78-2-22-28-16 | 6 | AABBDD | 41.6666667 | 39.44 | 15.2207327 | 3.49963846 | 5.72504631 |
| C538-1-78-2-22-28-17 | 6 | AABBDD | 28.6666667 | 38.1395349 | 14.8404766 | 3.39595859 | 5.86773535 |
| C538-1-78-2-22-28-18 | 6 | AABBDD | 39 | 40.8547009 | 15.6717079 | 3.46084551 | 6.02284991 |
| C538-1-78-2-22-28-2 | 6 | AABBDD | 46 | 39.9275362 | 15.4697161 | 3.53301031 | 5.85409734 |
| C538-1-78-2-22-28-3 | 6 | AABBDD | 39 | 41.2820513 | 15.4187017 | 3.51877656 | 5.8881548 |
| C538-1-78-2-22-28-4 | 6 | AABBDD | 42 | 40.952381 | 15.7534401 | 3.58185243 | 5.90149076 |
| C538-1-78-2-22-28-7 | 6 | AABBDD | 36.3333333 | 39.266055 | 15.2974573 | 3.46136475 | 5.84717069 |
| C538-1-78-2-22-28-9 | 6 | AABBDD | 39 | 38.8034188 | 15.4429805 | 3.46768075 | 5.93867804 |
| C538-1-78-2-22-33-21 | 6 | AABBDD | 55 | 35.2727273 | 14.4552638 | 3.33371731 | 5.76799846 |
| C538-1-78-2-22-33-23 | 6 | AABBDD | 48.3333333 | 39.0344828 | 15.1974746 | 3.48896911 | 5.83041155 |
| C538-1-78-2-22-33-25 | 6 | AABBDD | 42.3333333 | 40.1574803 | 15.4109646 | 3.47817209 | 5.86280659 |
| C538-1-78-2-22-33-37 | 6 | AABBDD | 56.6666667 | 40.8823529 | 15.8789938 | 3.5506625 | 5.97536223 |
| C538-1-78-2-22-62-19 | 6 | AABBDD | 40.6666667 | 39.5901639 | 15.3345447 | 3.49362636 | 5.86376468 |
| C538-1-78-2-22-62-4 | 6 | AABBDD | 40.3333333 | 39.4214876 | 15.5204869 | 3.4623829 | 5.98087373 |
